# A practical guide for generating unsupervised, spectrogram-based latent space representations of animal vocalizations

**DOI:** 10.1101/2021.12.16.472881

**Authors:** Mara Thomas, Frants H. Jensen, Baptiste Averly, Vlad Demartsev, Marta B. Manser, Tim Sainburg, Marie A. Roch, Ariana Strandburg-Peshkin

## Abstract

The manual detection, analysis, and classification of animal vocalizations in acoustic recordings is laborious and requires expert knowledge. Hence, there is a need for objective, generalizable methods that detect underlying patterns in these data, categorize sounds into distinct groups, and quantify similarities between them. Among all computational methods that have been proposed to accomplish this, neighborhood-based dimensionality reduction of spectrograms to produce a latent-space representation of calls stands out for its conceptual simplicity and effectiveness. Using a dataset of manually annotated meerkat (*Suricata suricatta*) vocalizations, we demonstrate how this method can be used to obtain meaningful latent space representations that reflect the established taxonomy of call types. We analyze strengths and weaknesses of the proposed approach, give recommendations for its usage and show application examples, such as the classification of ambiguous calls and the detection of mislabeled calls. All analyses are accompanied by example code to help researchers realize the potential of this method for the study of animal vocalizations.

## INTRODUCTION

Unsupervised dimensionality reduction projects data into low-dimensional space with the aim of visualizing underlying structure and aiding its detection. In contrast to supervised methods, it is not designed to separate known classes, but to reveal previously unknown structure and patterns in unlabeled datasets. Enabling exploration of high dimensional datasets in a purely data-driven way makes unsupervised dimensionality reduction applicable to many research fields, among them the study of animal vocalizations. While it is often known that there is some underlying structure in vocalization datasets (e.g. different types of calls), it can be difficult to categorize signals into distinct types or quantify similarities between them. If done manually, e.g. by human listeners, there is often disagreement between annotators, partly because of a lack of clear-cut rules for categorization and partly because many vocal repertoires are graded to some extent [1]. Computational methods that tackle these challenges in a more objective and quantifiable way are thus highly desirable. Unsupervised dimensionality reduction can provide the basis for such computational tools by projecting entire datasets of vocalizations to 2D or 3D space, thus allowing one to visualize underlying structure and facilitating the identification of clusters of highly similar signals (i.e. call types). Such methods have been extensively used to study vocalizations, but have mostly been applied to acoustic feature vectors, a type of encoding where vocalizations are described by parameters such as their fundamental frequency, mean spectral entropy, bandwidth or cepstral peak prominence (reviewed in [2]). Disadvantages of this approach are that the resulting visualization in 2D or 3D space varies greatly with the choice of acoustic features and that feature extraction is often not trivial and requires expert knowledge. Recently, Sainburg et al. [3] proposed a method to generate meaningful latent space representations of animal vocalizations that works directly on spectrograms, thus holding the promise of providing a less biased, more objective, and easier to implement method to study vocal repertoires within and across species. Using Uniform Manifold Approximation and Projection (UMAP), they directly mapped spectrograms into low dimensional latent space. In addition to the identification of call type groups through clustering in latent space, they showed that the low dimensional representations can be used to study vocal encoding of individual identity, make cross-species comparisons and analyze sequential organization of vocalizations. To make this simple yet effective approach more accessible, we provide a tutorial for the generation of such representations using a dataset of meerkat (*Suricata suricatta*) calls as an example. The meerkat repertoire is an ideal example use case as it has been extensively studied [4], yet holds many of the challenges that are typical in the field of bioacoustics: There are a number of distinct and well-characterized call types, but also some degree of gradation, with calls falling in between those types. There is disagreement between human labellers on correct categorization of these types and there are sub-types which have not yet been fully described. By comparing the patterns resulting from unsupervised dimensionality reduction to the manual categorization of calls by human expert labellers, we show strengths and weaknesses of the UMAP approach by Sainburg et al. [3]. In addition, we discuss the choice of various preprocessing steps and dimensionality reduction hyperparameters and provide recommendations as well as application examples for researchers who wish to apply this method to their own data.

## EXPLANATION OF THE METHOD

The approach of Sainburg et al. [3] is based on two core concepts: First, the encoding of animal vocalizations as row-wise concatenated spectrograms as opposed to vectors of acoustic features. Second, the unsupervised dimensionality reduction with UMAP, a method that is based on manifold learning and topological data analysis.

Each animal vocalization is first transformed into a spectrogram, a visual representation of the frequency content of the signal over time. Then, all rows of the spectrogram are concatenated to generate a long, numerical vector [3]. Hence, each vocalization is encoded as a vector (or high-dimensional datapoint), where each field (or dimension) contains one “pixel” of the original spectrogram, i.e. the signal magnitude at a certain point in frequency and time. Computing the Euclidean (or other) distance between these spectrogram vectors can then be used as a measure of acoustic similarity, which is essentially based on the element-wise comparison of spectrograms (similar to spectrogram cross-correlation [5]). UMAP is then used to visualize the structure of the data in a low-dimensional space (e.g. 2D, 3D) as well as to facilitate the detection of clusters of similar vocalizations.

UMAP was first described in 2018 as a novel dimensionality reduction method that produced results similar to t-Stochastic Neighbor Embedding (t-SNE) [6], but is based on a different mathematical framework and is computationally faster and more scalable [7]. In brief, UMAP constructs a weighted neighborhood graph from high-dimensional data and finds a lower dimensional representation with similar topological properties through a stochastic learning process. This general objective makes UMAP very similar to other neighborhood graph based algorithms (e.g. t-SNE [6] or LargeVis [8]), and differentiates it from algorithms that, for example, aim to preserve all pairwise distances (e.g. Sammon mapping [9], Multi-Dimensional Scaling [10]) or variation in the data (e.g. Principal Component Analysis (PCA) [11]). The core idea of UMAP and other neighborhood graph methods is to emphasize the preservation of local over global structure in the inherently lossy process of dimensionality reduction (as it is generally impossible to maintain the exact distance structure of high-dimensional data in lower dimensional space). Hence, given a dataset of different types of animal vocalizations, an embedding generated with UMAP (or t-SNE, LargeVis etc.) will more accurately reflect the closeness of similar vocalizations in space, while distances between more dissimilar vocalization types will not necessarily resemble those in the original space. Further, relative local density of the data is not preserved, meaning that dense or more loose point clouds in latent space do not necessarily reflect density of the original data (however a density-preserving version for UMAP has been recently been published [12]). While a detailed discussion of the UMAP algorithm is beyond the scope of this manuscript, it is important to keep its basic properties in mind when interpreting embeddings, as their consequences could otherwise be misinterpreted ([7] and [13] recommended for further reading).

## THINGS TO CONSIDER BEFORE USING THIS METHOD

### Input requirements

The approach requires a dataset of sound files, each containing a single vocalization or syllable as input. Note that it may be necessary to extract such vocalizations from acoustic recordings in a preprocessing step which is not covered here. (We recommend dynamic threshold segmentation as proposed in [3] or other acoustic event detection, see [14] for an overview). Ideally, the start and the end of the sound file correspond to start and end of the vocalization. If there are delays in the onset of the vocalizations, these should be the same for all sound files. Otherwise, varying onsets may make these vocalizations appear dissimilar or distant in latent space. If it is not possible to mark the start times correctly, the pipeline can be adapted, e.g. to allow for time-shifting (see supplementary information).

### Considerations of sample size and constitution

While there is no definite minimum sample size for UMAP, it is not recommended to use UMAP on datasets with less than 100 samples in total. For our dataset, N=50 vocalizations of each type (N=350 in total) were sufficient to achieve the same degree of call type clustering as with any higher number of samples (see supplementary information P3). Strong over- or underrepresentation of specific vocalizations in the dataset (e.g. class imbalance) was also unproblematic for our dataset (see supplementary information P2). However, it may be advisable to downsample heavily overrepresented types to improve the readability of the visualizations.

### Constraints

The biological meaningfulness of distances in latent space, i.e. how well they reflect similarity of vocalizations, depends mostly on the parameters for spectrogram generation and transformation and the distance metric selected for UMAP. Therefore, these must be chosen with care and adapted for each dataset. We provide some recommendations for generating, denoising and transforming spectrograms and compare different distance metrics in sections “Caveats and pitfalls” and supplementary information P7. Lastly, it is important to keep in mind that distances in UMAP space do not faithfully reflect the distances in original space, as UMAP is designed to favor the preservation of local over global structure.

## WORKED EXAMPLES

We present all steps of the computational pipeline as proposed by Sainburg et al. [3] using a dataset of N=6,428 meerkat calls as an example (see supplementary information P1 for a detailed description of data collection and cleaning). The vocalizations in this dataset were between 50 – 500 ms long and had been manually labelled as one of seven call types: aggression (*agg*), alarm (*al*), close call (*cc*), lead (*ld*), move (*mo*), short note (*sn*) or social call (*soc*). Very noisy calls and ambiguous calls had been previously removed from the dataset (supplementary information P1). All computations were performed with Python 3.8.

### Generation of spectrograms

First, a spectrogram is constructed from each audio file via a series of discrete short-time Fourier Transforms (STFT). Audio files contain time-series data (sound pressure over time) and the sampling rate (*sr)*, which indicates how many audio samples were acquired per second (Hz). STFT divides this data into chunks, decomposes each chunk into a vector of sound magnitude per frequency bin through Fourier transformation and puts the vectors (or FFT frames) side-by-side. The resulting spectrogram is a two-dimensional matrix *M*, where *M[i,j]* denotes the magnitude of the signal for a given frequency interval *i* at time point *j*. We used the STFT function from *librosa v0*.*8*.*0* [15], but other packages provide the same functionality. Several hyperparameters define the spectrogram’s resolution in time and frequency and prevent the occurrence of artifacts: The parameter *n_fft* determines how many audio datapoints go into one FFT frame and thus the frequency resolution of the spectrogram (frequency resolution in Hz = *sr*/*n_fft*). The parameter *hop_length* determines how many audio samples lie between adjacent FFT frames and is often set smaller than *n_fft*, so that there is overlap between adjacent frames of the spectrogram. This improves the odds of FFT frames falling near the boundary of changes in the signal, and thus improves the visibility of signals. In addition, a window function is applied to the audio data of each FFT frame to prevent spectral leakage [16]. For our dataset, we set *n_fft* such that 30 ms of audio data were used to generate one time frame (and a same-sized Hann window function) and set the hop length such that 3.75 ms passed between adjacent FFT frames, resulting in an overlap of 87.5% between successive frames. We defined these parameters in seconds and calculated *n_fft* and *hop_length* for each audio file based on sampling frequency to ensure consistent temporal resolution of audio files (our dataset contained files with *sr* = 48,000 and *sr* = 8,000 Hz). As the highest frequency that can be detected without aliasing is at the Nyquist rate (one-half of the sampling rate), spectrograms of samples with higher sampling rates will have a larger frequency range than those with lower sampling rates (24,000 Hz vs. 4,000 Hz). Therefore, we only used frequency bins between 0-4000 Hz across all spectrograms.

### Preprocessing of spectrograms

The spectrograms then undergo two modifications in order to emphasize biologically relevant features: (1) The frequency bins (Hz) are transformed to Mel bins based on the Mel-scale, a logarithmic, experimentally determined psycho-acoustic pitch scale [17]. Mel-transformation emphasizes differences between perceptually distinct calls by distorting the frequency axis to match the non-linear hearing abilities of humans. Depending on the study species of interest and their hearing abilities, it may not always be advisable to apply. We transformed all spectrograms using a Mel filterbank of 40 coefficients between 0 - 4,000 Hz. (2) The energy content of the spectrogram is then transformed to a Decibel scale to reflect how the human auditory system perceives loudness logarithmically [18]. As we used the maximal power of the spectrogram as reference, this step also provides a normalization for varying loudness of the audio files. Note that this may also be undesirable if you wish to distinguish vocalizations based on their loudness.

### Generating input vectors for UMAP

Next, each spectrogram is *z*-transformed to normalize for differences in overall intensity between calls, and padded with zeros up to the maximal call duration in the dataset (500 ms), so that all spectrograms in the dataset are of equal length (UMAP requires a static number of attributes). All spectrograms are then row-wise concatenated to generate feature vectors (spectrogram-vectors), which can be conceptualized as points in high-dimensional space.

## UMAP

Spectrogram-vectors are mapped into low-dimensional space (2D and 3D) using UMAP from *umap-learn* [7]. UMAP builds an approximate nearest neighbor graph from the datapoints in original space (here, spectrograms of vocalizations) by computing a user-defined distance between input vectors (default: Euclidean) and then finds a low-dimensional representation that preserves the structure of the graph in an iterative optimization procedure. Even though many properties of the UMAP algorithm can be specified, the default values (with the exception of the parameter *min_dist*=0, which is recommended for clustering) provided good results for our dataset and were also proposed in Sainburg et al. [3] (see supplementary information P7 for a more detailed analysis of the effects of different UMAP hyperparameters). Projecting the calls into 2D and 3D space (*n_components*=2 or *n_components*=3) and coloring the datapoints by their manual labels revealed structures with few distinct clusters (Figure 1), but clear separation of the manually annotated call types.

**Figure 1:**
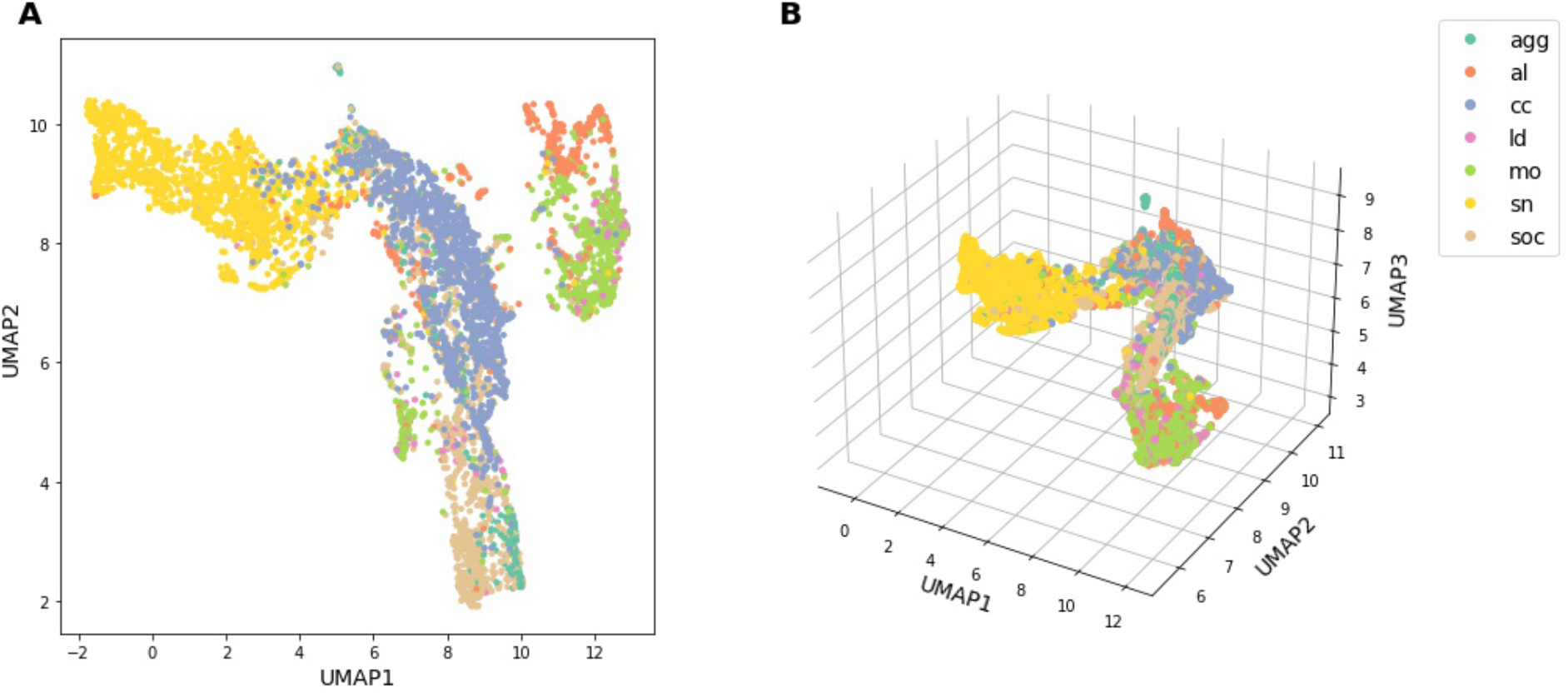
Latent space representations of meerkat vocalizations in A) 2D and B) 3D, color-coded by manual call type labels.

### Interactive visualization

To explore the 3D latent space representation in more detail, we developed an interactive visualization tool with audio playback (Figure 2, demonstration video and code tutorial in the provided code repository). Hovering over datapoints triggers the display of the respective spectrogram next to the plot, as well as a table containing metadata of the datapoint (e.g. meerkat identifier, sex and social status for our dataset), while a mouse-click on the datapoint triggers the audio playback of the respective call.

**Figure 2:**
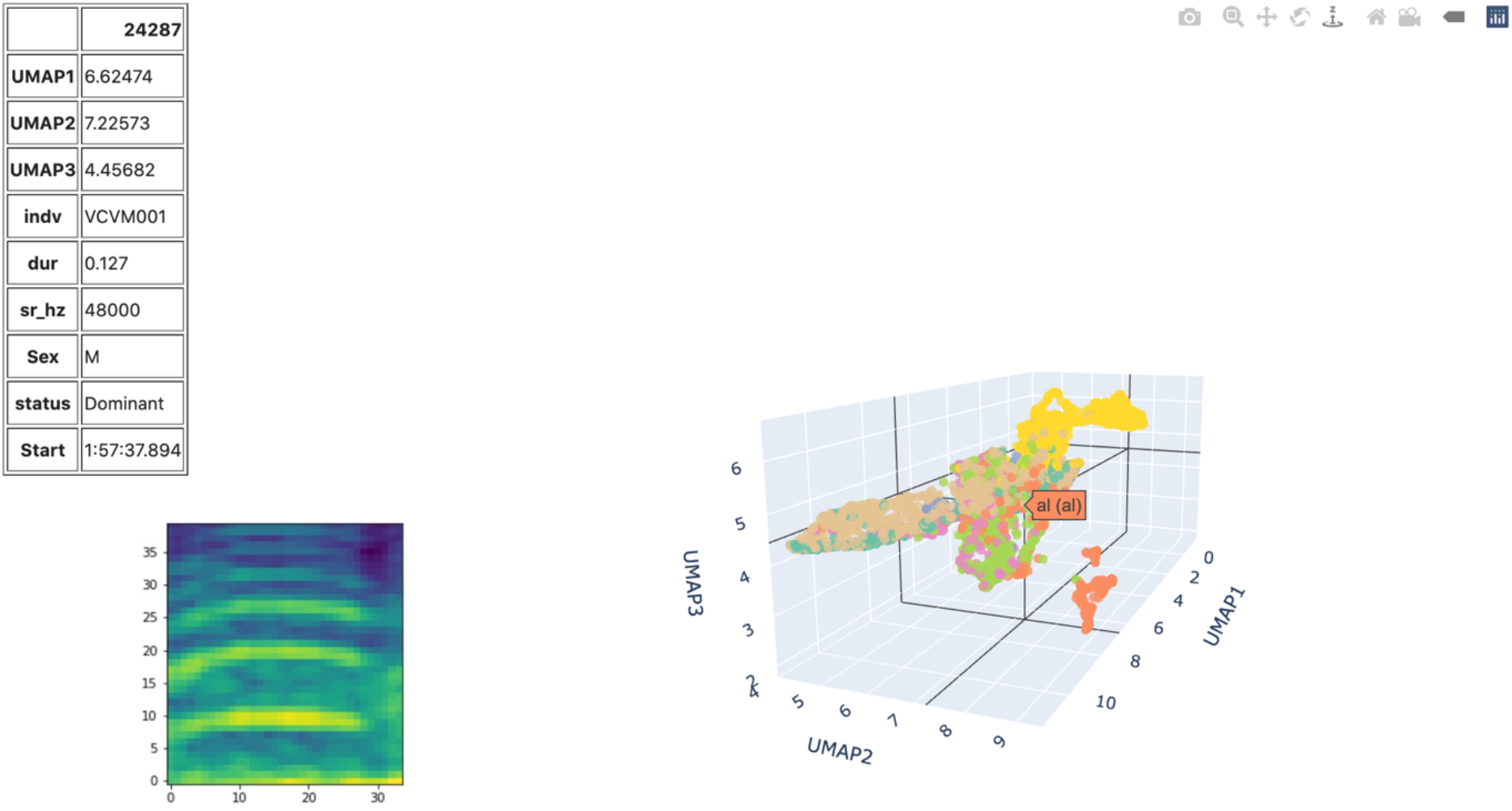
Screenshot of the interactive visualization tool, demonstrated with meerkat dataset (N=6,428). Table on the left indicates the identifier of the meerkat (indv), call duration in s (dur), samplerate (sr_hz), sex and social status of the individual. The spectrogram of the respective call is displayed below. The plot on the right shows the representations in 3D UMAP space, colored by manual call type label.

### Evaluation of the latent space representations

While it is clear that a good representation is one where similar vocalizations are close together and dissimilar ones are distant, the development of metrics and methods for evaluating embedding quality is an open problem in the field of dimensionality reduction and the choice of embedding quality metrics is largely dependent on the experimenter’s goals in embedding. Here, we discuss a set of embedding metrics that we consider relevant to the evaluation of vocal repertoire embeddings, both for completely unlabeled and partially labeled datasets.

If no information on call types is available, the association of distance in latent space with call type similarity can only be qualitatively assessed, e.g. by exploring the space through the interactive visualization or by randomly pulling out example calls and their nearest neighbors. If some or all vocalizations in the dataset are labelled (as in our meerkat dataset), the clustering of call type groups in latent space can be quantitatively assessed for the labelled data points and used as a proxy of how well the embedding reflects call similarity (assuming that calls of the same type are more similar than those of different types). Possible metrics are the Silhouette coefficient applied to the manual call type groups, the average percentage of calls of the same type in the local neighborhood or the adjusted Rand index to compare the partition obtained from unsupervised clustering to the manual labels. In this tutorial, we also apply these metrics to the original, high-dimensional space to demonstrate the effects of dimensionality reduction. Lastly, structure preservation (e.g. the performance of UMAP) can be evaluated by assessing the nearest neighbor preservation and the correlation of distances in low- and high-dimensional space (see supplementary information P5, P6). In general, it is important to keep in mind that the quality of the embedding depends on both i) how well the distance metric in the original space reflects similarity between vocalizations and ii) how well the dimensionality reduction has preserved the structure of the data.

#### Random nearest neighbor visualization

To assess whether a call’s nearest neighbors are indeed acoustically similar, we randomly select calls from the dataset along with their k nearest neighbors, display the spectrograms, visually assess their similarity (Figure 3) and/or play back the audio (see “Interactive visualization”).

**Figure 3:**
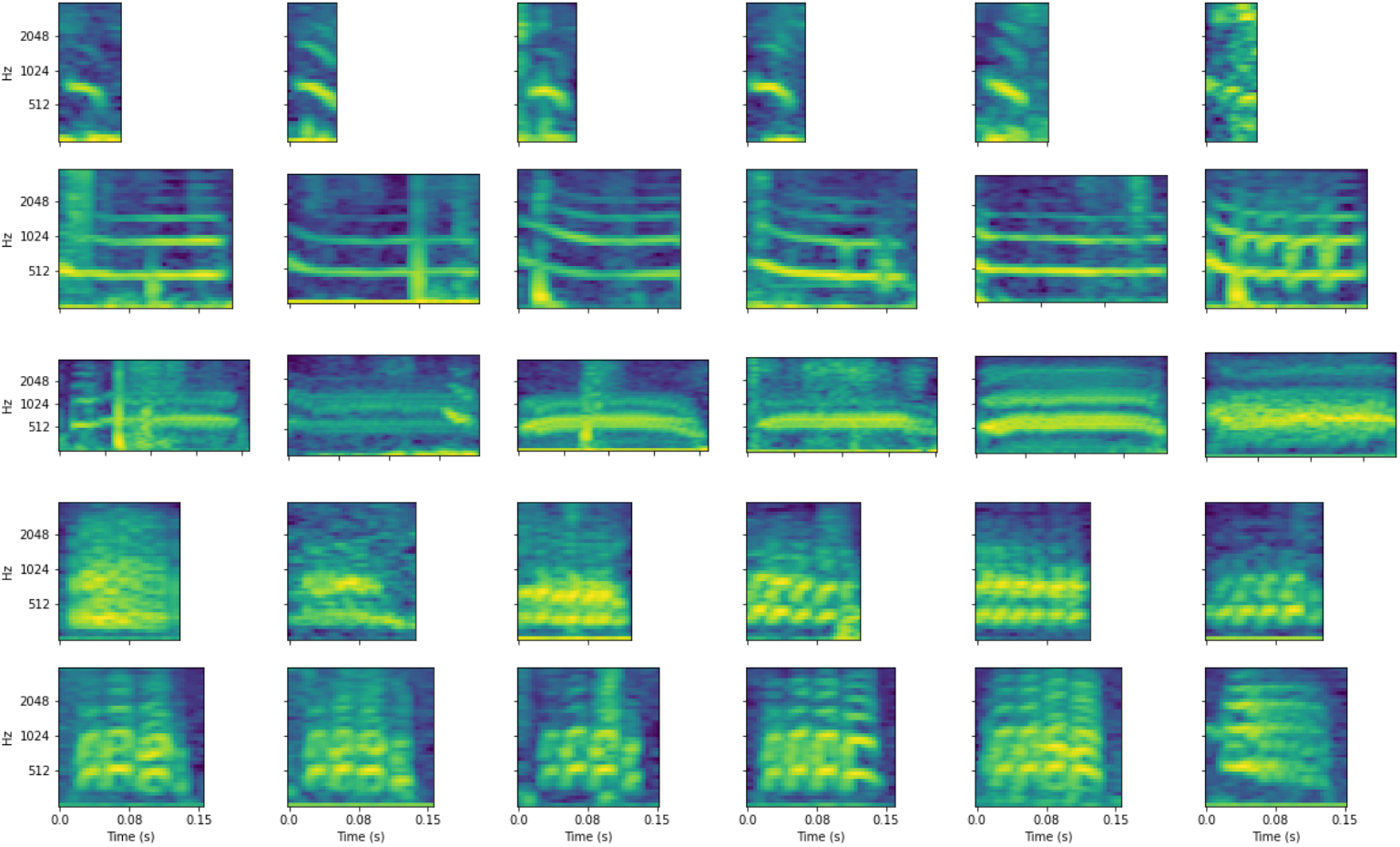
Spectrograms of five randomly selected calls from the dataset (first column) and their k=5 nearest neighbors in latent space (columns 2-6).

#### Nearest Neighbor Metrics

To evaluate to what degree acoustically similar calls (e.g. with the same manual label) cluster in latent space, we assess the probability that a call is surrounded by calls of the same type in latent space. (This can also be performed for a labelled subset of the full dataset.) For a given call type label *i*, we select all calls of that label, identify their k nearest neighbors in latent space and note their type. We then analyze the composition of these neighbor labels (e.g. 10% of the labels are close call, 23% lead call etc.) and use the observed frequency as an estimate for the probability *P* of encountering calls of this particular type among the k nearest neighbors of calls of label *i*.

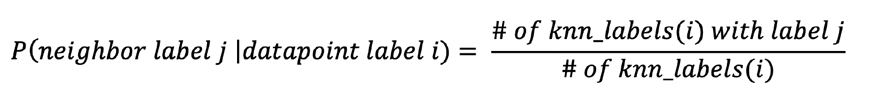

Here, *knn_labels*(*i*) is the list of labels of the k nearest neighbors of all datapoints with label *i*.

Doing this for all classes (i.e. manually labelled call types), we obtain a square evaluation matrix where each field [i,j] represents the probability *P* (expressed in %) for a call of type *i* to have a neighbor of type *j* (Figure 5A and 5B).

**Figure 5:**
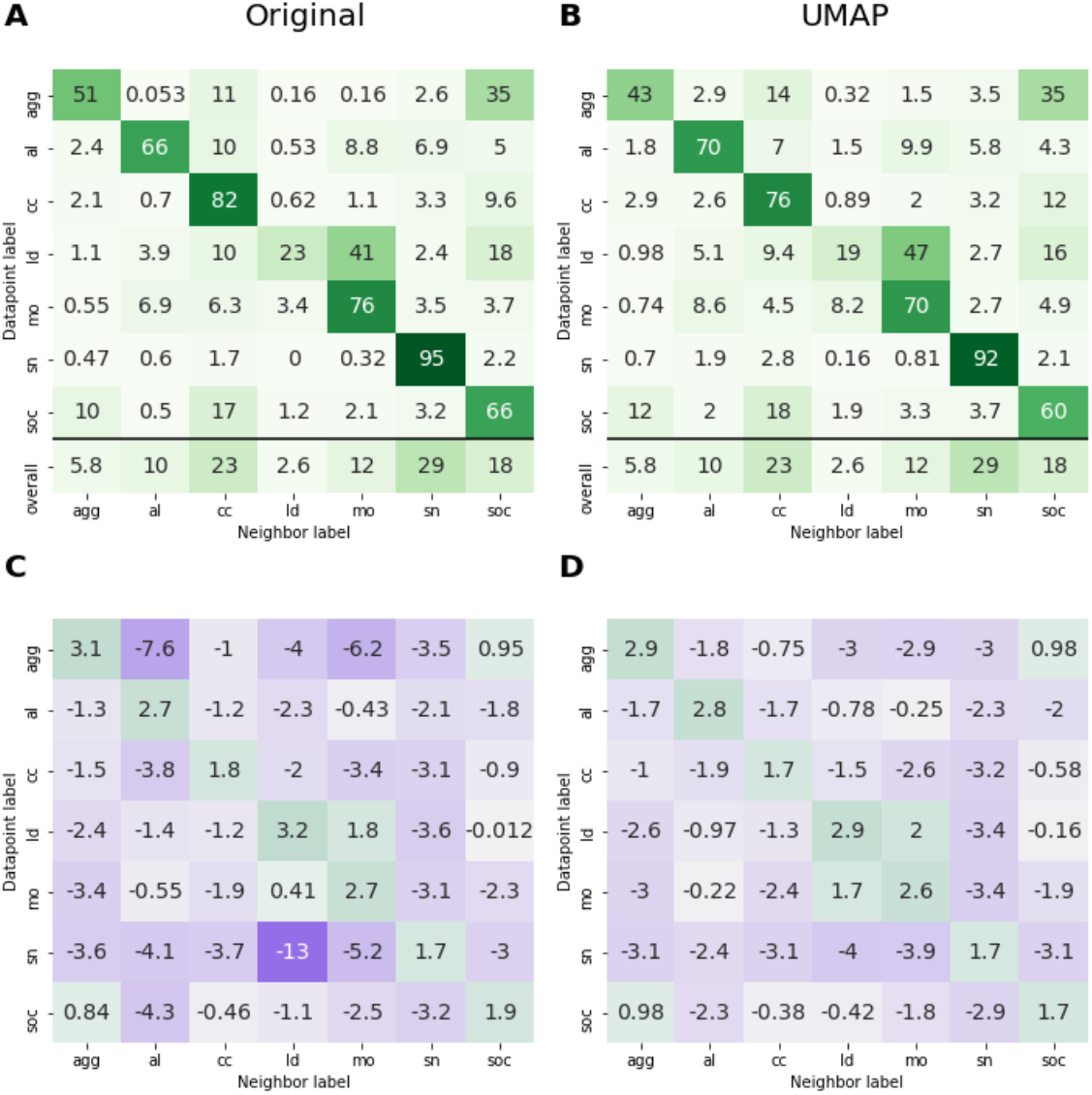
Evaluation matrix in original (A, C) and 3D UMAP (B, D) space. A)-B) show average frequency (%) of datapoints with label x among the nearest neighbors of a datapoint with label y. For ease of interpretation, we added a row for the random chance expectation of encountering a neighbor with this label in the dataset (“overall”, e.g. frequency of this call type in the dataset). C)-D) show log2-transformed ratio of observed frequency vs. frequency expected by chance (e.g. class frequency). Colors are mapped on a violet-green scale from minimum to maximum value.

Since this probability is not normalized to varying call type frequencies in the dataset (i.e. it is more likely to have a common call type in the neighborhood than a rare one by chance alone), we divide it by the probability of encountering calls of this type by random chance alone (i.e. the frequency of this label in the dataset). The resulting score can then be interpreted as the fold increase or decrease in likelihood of observing this many neighbors over the random chance expectation. To make the score symmetric around zero, we apply a log2 transformation and obtain the normalized score *P*_*norm*_ (Figure 5C and 5D).

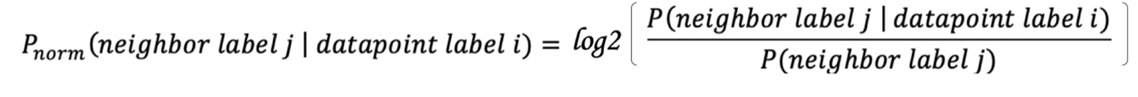

with *P(neighbor label j)* being the probability of observing a neighbor with label *j* due to random chance alone (e.g. the frequency of label j in the dataset).

To capture the quality of an embedding in a single score, we calculate the unweighted average of *P* (or *P*_*norm*_) for same-class neighbors over all classes (the diagonal of the evaluation matrix), thus obtaining the summarized score *S* (or *S*_*norm*_). We explicitly chose the unweighted average, so that the same-class neighbor probabilities of each call type have equal weight in the final score and the final score is not biased towards the same-class probabilities of the more frequent call types in the dataset. When comparing different embeddings of the same dataset, i.e. with the same distributions of call type frequencies, we report the unnormalized score *S*, which can be interpreted as the average frequency of same-class labels among the k nearest neighbors of each call type.

For our meerkat dataset, the summarized score *S* was 61.3%, e.g. for any given call type, on average 61.3% of the k=5 nearest neighbors were of the same type. This indicates that calls of the same type were found much more often in close neighborhoods than expected by random chance alone (random chance expectation: 14.7%). The percentage of same-class neighbors varied among call types, with the highest score for *sn* calls (92%), which was also the most frequent call type in the dataset (29%). When normalized to the random chance expectation, *ld* and *agg* calls had the highest normalized probability of same-class neighbors (*P*_*norm*_=2.9, i.e. frequency of same-class neighbors was 7.5-fold (2^2.9^) higher than expected by random chance) (Figure 5D). The normalized neighbor metrics indicate that move and lead calls, as well as social and aggressive calls, are often found in close vicinity of each other and these calls are indeed acoustically (and functionally) similar (see supplementary Figure S1). Altogether, we recommend inspecting the full evaluation matrices of both scores to get a comprehensive overview of the neighborhood probabilities of different vocalization types.

To understand the effects of UMAP, we calculated the same evaluation matrices in original, high-dimensional space (i.e. spectrogram vector space). While the overall patterns between UMAP and original space were very similar, the original space had slightly higher quality scores (*S*=65.59 versus *S*=62.81 and *S*_*norm*_=2.45 versus *S*_*norm*_=2.4), indicating that the local neighborhood (k=5) was not exactly preserved in UMAP. When re-calculating the quality score *S* for a different number of k nearest neighbors and thus investigating a more global neighborhood, we found that within a larger neighborhood (k>25), *S* was higher in UMAP than in original space (Figure 6). This shows that while UMAP does not accurately preserve the closest nearest neighbors, it does improve the overall clustering of similar datapoints.

**Figure 6:**
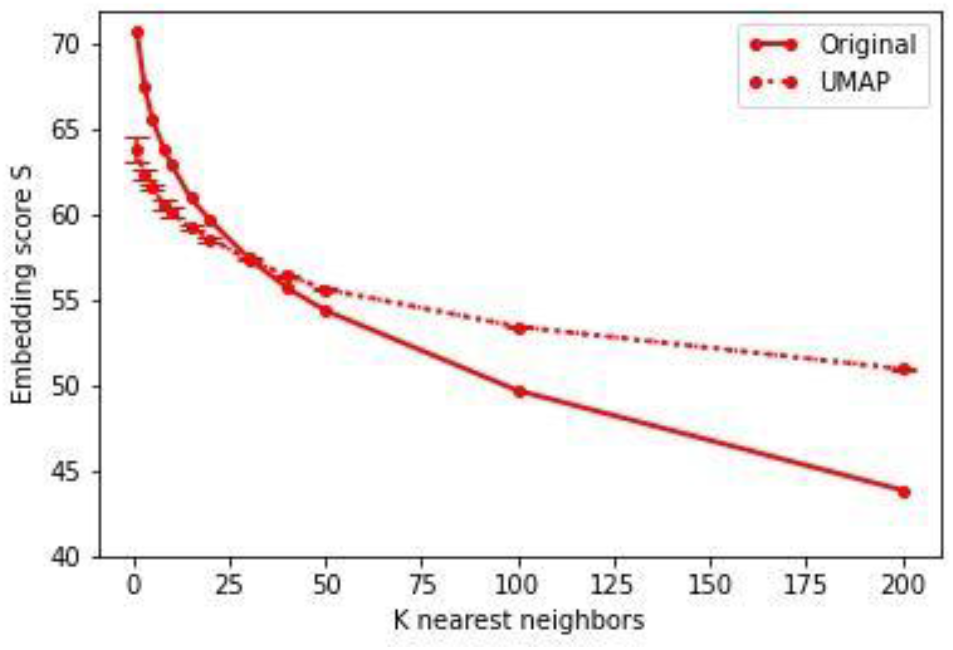
Comparison of embedding score S for different k nearest neighbors in original vs. 3D UMAP space. UMAP line represents mean and standard deviation of n=5 UMAP runs.

#### Within-vs. between call type distances

We also investigated the distribution of pairwise distances within a call type group vs. between calls of a different type. Again, we show results for original and UMAP space to demonstrate the effects of UMAP.

The average distance to datapoints of the same type was smaller than the average distance to datapoints of a different type for all call types in UMAP, but not in original space (Figure 7). This illustrates the effectiveness of UMAP in generating tighter clusters of similar datapoints, which facilitates the detection of patterns and structure in the data. In our dataset, the best separation of within-vs. between-call type distances was obtained for *mo* and *sn* calls. *Cc* and *al* calls were less separated from other call types, even though they had similarly high same-class nearest neighbor frequencies (*mo* 70%, *sn* 92%, *al* 70%, *cc* 76%), illustrating the different types of class separation that are captured by within-vs. between-class distances as opposed to the nearest neighbor metrics.

**Figure 7:**
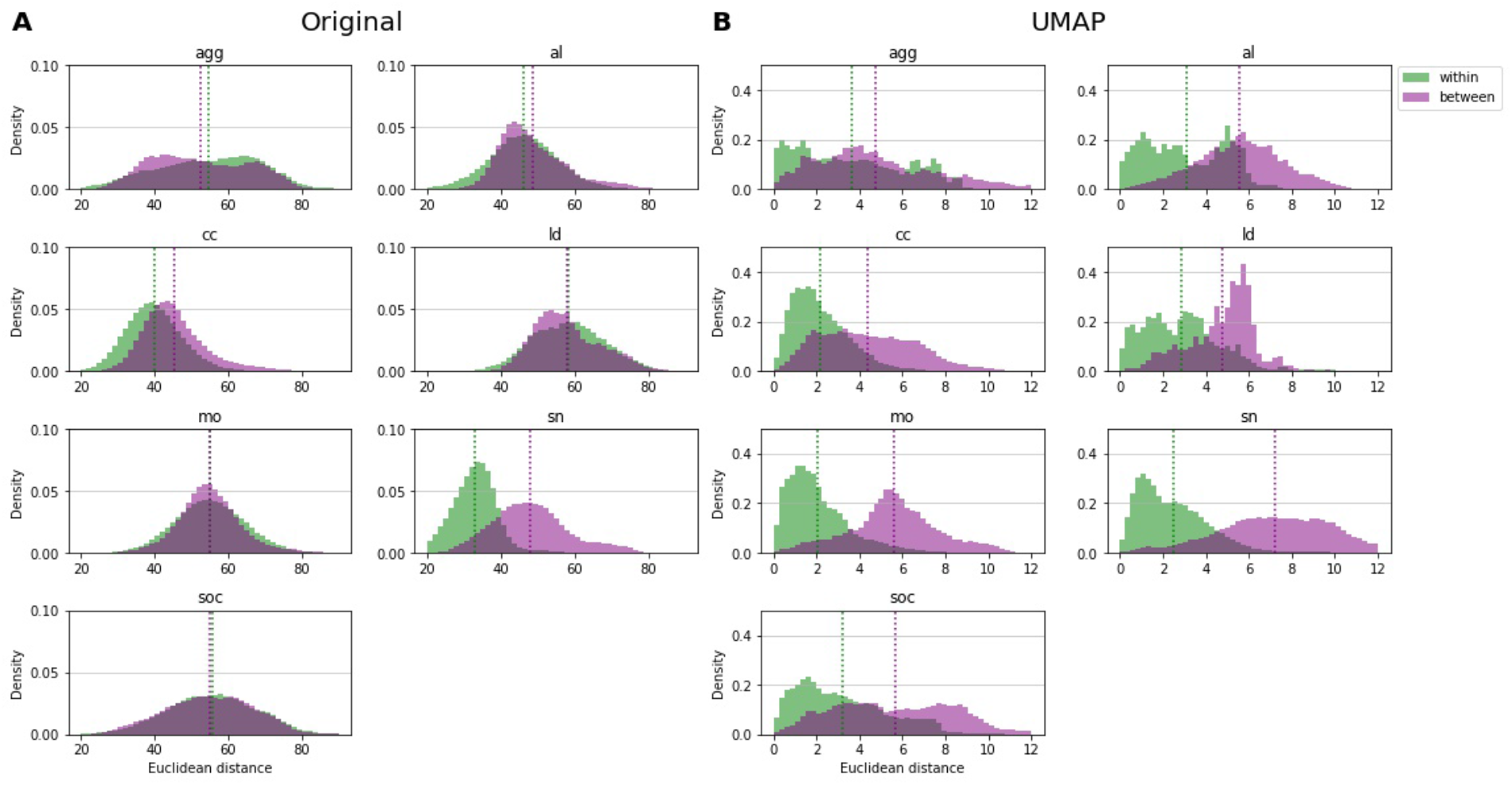
Distribution of pairwise distances between datapoints of the same call type (within) vs. between datapoints of different call types (between), shown for each call type separately and for A) original space and B) 3D UMAP space. Dotted, vertical lines show the mean. Note the different x-and y-scales for original vs. UMAP space.

#### Silhouette plot

As another means to quantify the global clustering by call type, we calculated the silhouette values of the manual label clusters in original and UMAP space using the implementation of *scikit-learn* [19] and plotted the scores for all datapoints sorted by call type group (Figure 8). The silhouette value indicates how close datapoints are to their own cluster compared to other clusters and is defined as:

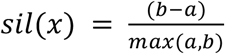

with *a* being the mean intra-cluster distance in the cluster of datapoint x and *b* the mean distance between x and the nearest neighboring cluster. A positive score thus indicates that this datapoint is near elements of the same cluster, whereas a negative score indicates that it is closer to elements of another cluster.

**Figure 8:**
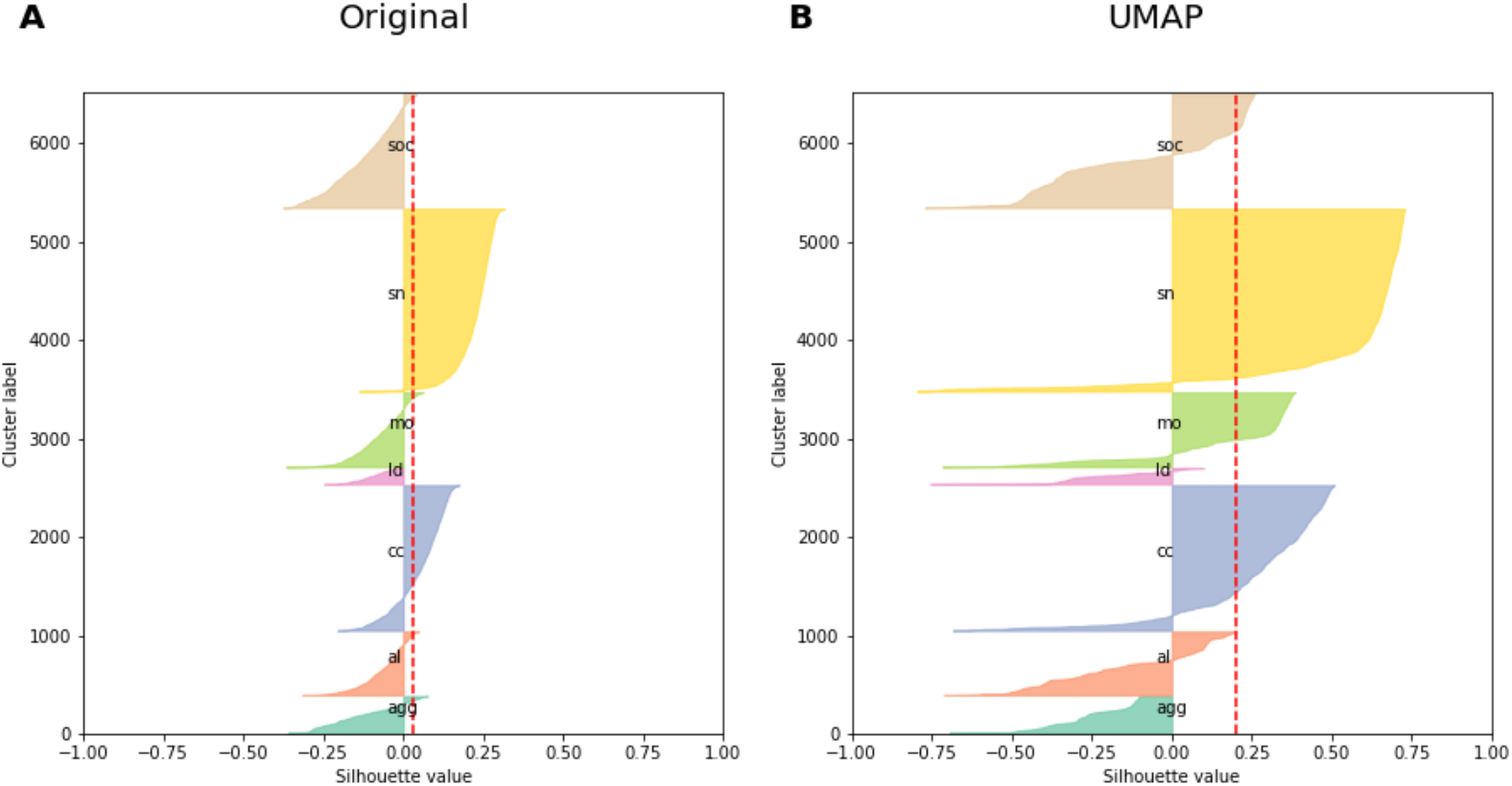
Silhouette plots for manual label clusters in A) original and B) UMAP space. Datapoints are sorted by label and silhouette values are displayed for each datapoint. Dotted red line indicates the average for all datapoints (SIL).

The comparison of the average silhouette value (= Silhouette coefficient, SIL) of manual label clusters in UMAP versus original space also confirms that dimensionality reduction improved the global clustering of call types in space (SIL=0.03 for original, SIL=0.20 for UMAP) (Figure 8). However, the scores are low in both original and UMAP space, indicating that when looking at overall distances, the call types are not tightly clustered in any of the spaces. These low values are also in line with the visual impression of few distinct clusters in the meerkat vocal repertoire (Figure 2).

#### Evaluation of clustering

Another way to assess the quality of an embedding of a fully or partially labelled dataset is to perform clustering on the latent space and evaluate the agreement between the clustering results and the manually annotated call type groups. However, the results will not only depend on the quality of the embedding, but also on the performance of the clustering algorithm. While a detailed discussion of clustering algorithms is beyond the scope of this tutorial, it should be noted that clustering results can be spurious and unstable depending on the choice of algorithm and the structure of the data (e.g. when forcing clustering algorithms on homogenous data). Here, we present the results of Hierarchical Density-Based Spatial Clustering of Applications with Noise (HDBSCAN), as proposed in [3]. HDBSCAN returns high confidence clusters at the cost of leaving some calls uncategorized (“noise”), thus ensuring that calls of the same cluster are highly similar. We applied *hdbscan* [20] on the latent space representations using the default parameters, except *cluster_selection_method*=‘leaf’ (to obtain more fine-grained clusters) and *min_cluster_size*=64 (1% of the dataset), following the recommendations of Sainburg et al. [3]. We obtained 14 clusters in the meerkat dataset, while 48.8% of the data was categorized as noise (Figure 9).

**Figure 9:**
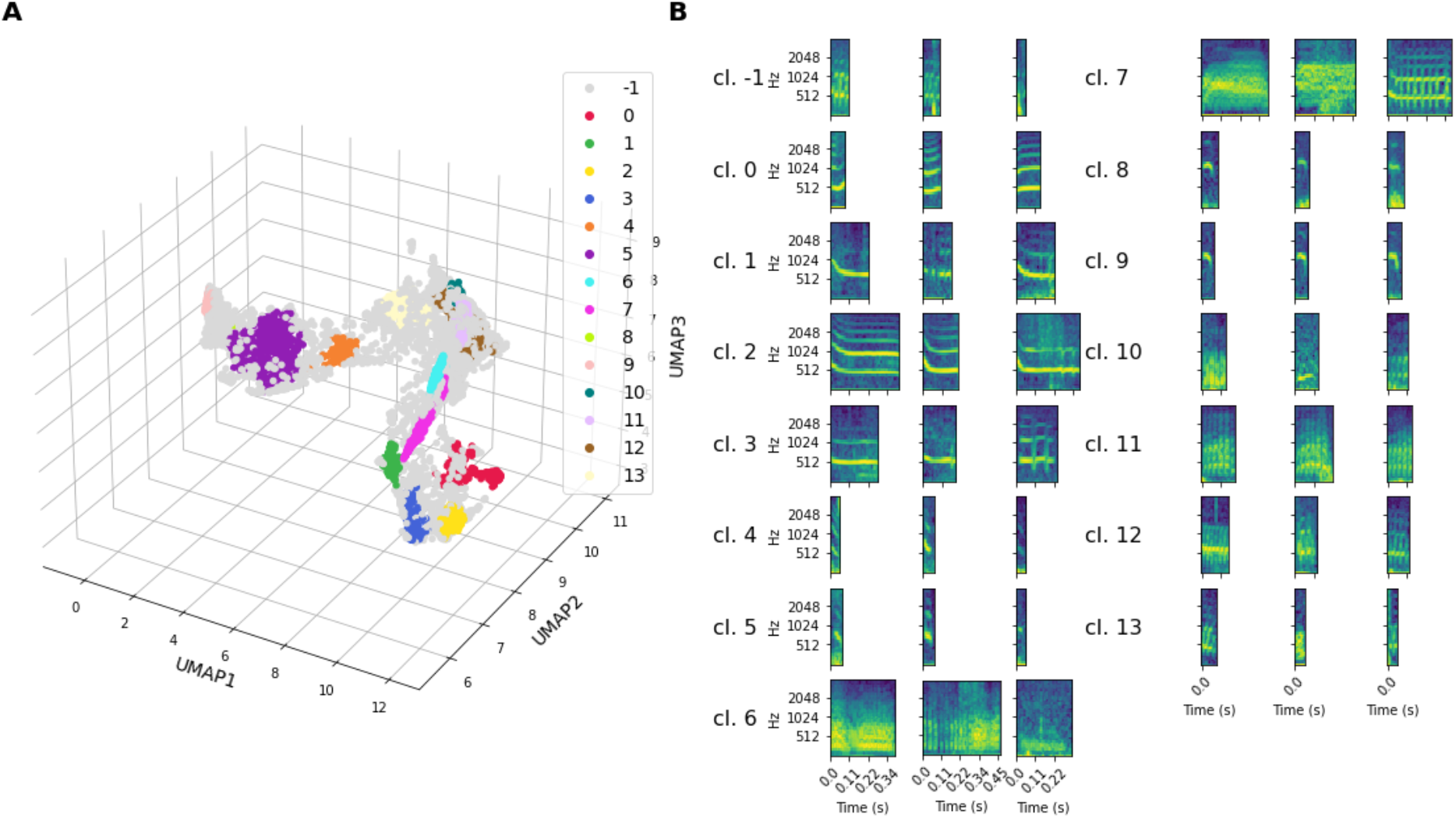
HDBSCAN clustering and n=3 randomly selected example calls of the different clusters. Cl. -1 contains noise.

We then assessed the agreement between the clusters obtained from HDBSCAN and the manual call type labels using adjusted (ARI) and unadjusted Rand index (RI) (*scikit-learn* [19]). We chose ARI and RI for their ease of interpretation, but alternative metrics for comparing partitions (e.g. Adjusted Mutual Information) are equally suitable. Comparing two partitions X and Y, the RI is is defined as:

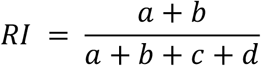

with *a* the number of pairs of datapoints that are in the same subset in X and the same subset in Y, *b* the number of pairs of datapoints that are in different subsets in X and different subsets in Y, and *c* and *d* the number of pairs of datapoints that are once in the same and once in a different subset in the two partitions X and Y. Hence, 100% agreement between two partitions results in RI=1. As even two random clusterings would have RI>0 due to chance alone, it is recommended to use ARI, the corrected-for-chance version of RI, where ARI=0 means that the agreement is no better than the random chance expectation.

After removal of the datapoints categorized as noise, our HDBSCAN clustering of meerkat calls had ARI=0.47 and RI=0.86, i.e. 86% of all pairs of datapoints were either in the same cluster and also had the same manual label or were in different clusters and also had different manual labels.

The composition of HDBSCAN clusters can be further investigated based on any available variable of interest (e.g. individual identity, sex, clan membership). We used the manual call type label for our dataset and found that nine of the HDBSCAN clusters were composed of a prominent call type fraction (>80%), and the remaining five contained a mix, with usually 2-3 prominent fractions (Figure 10C). Some of these mixed clusters contained call types that are acoustically similar (e.g. cl 1. *mo* and *ld* calls, cl. 6 *soc* and *agg* calls), whereas others contained call types that appear acoustically and visually (based on spectrograms) distinct (e.g. cl. 10 *al, cc* and *agg* calls). Closer inspection however showed that some mixed clusters did indeed contain similar calls. For example, the *al* calls in cl. 10 were mainly a specific subtype of *al* calls (terrestrial alarms), which are indeed acoustically more similar to *cc* and *agg* calls (with which they were clustered) than to the more common aerial alarm calls in the almost-pure alarm call cluster cl. 0 (Figure 11).

**Figure 10:**
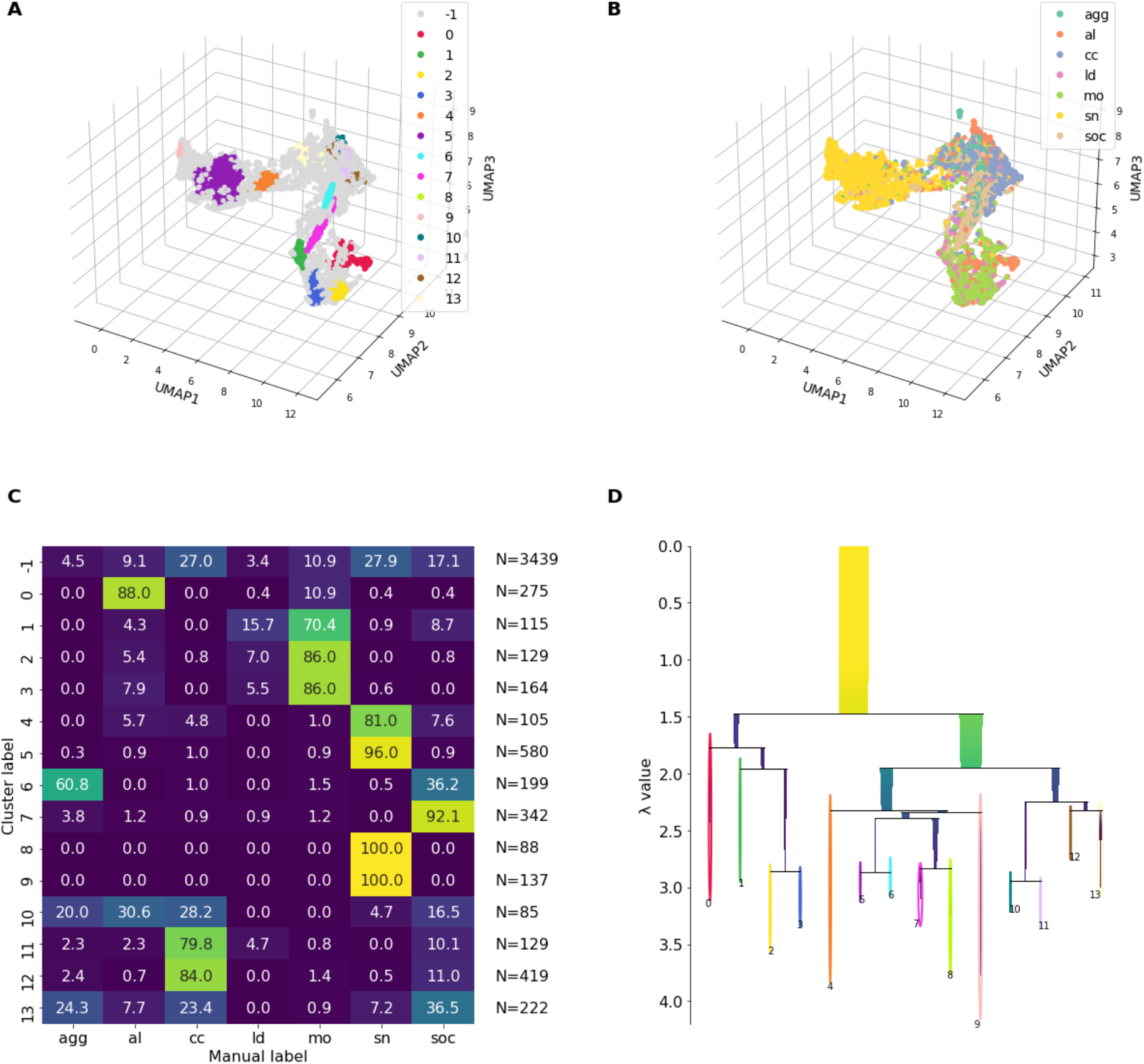
HDBSCAN clusters of the 3D UMAP embedded meerkat calls (A) as compared to the manual labels (B). C) shows composition of HDBSCAN clusters, i.e. the percentage of manual call types present in each cluster (row-wise percentages). Total numbers of calls in each cluster are reported as well. D) shows hierarchy of HDBSCAN clusters.

**Figure 11:**
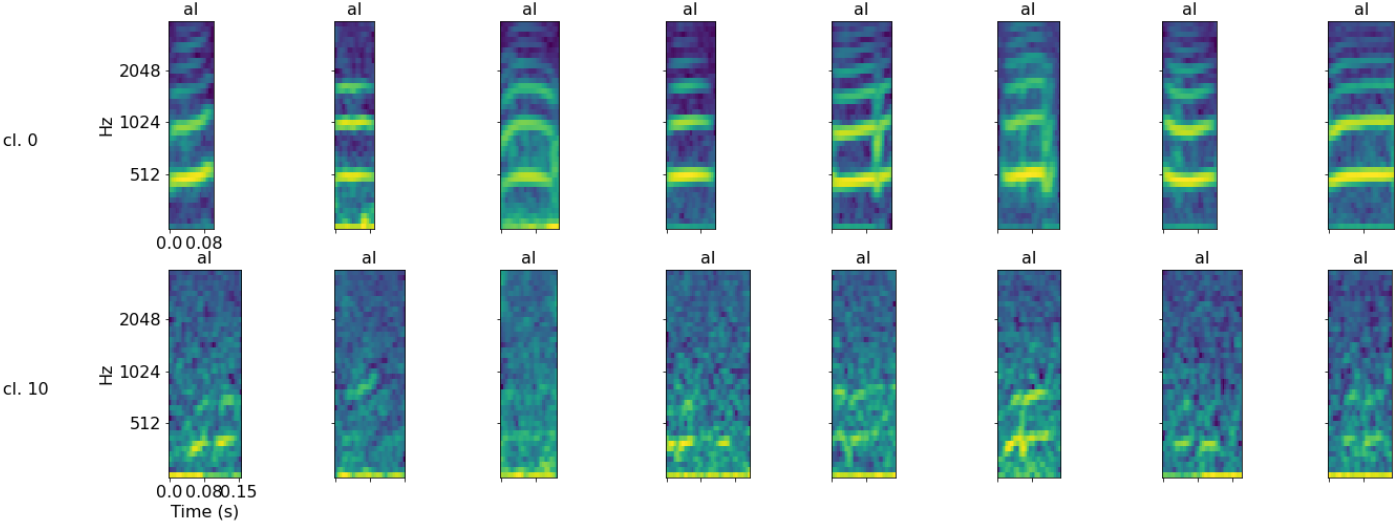
Example spectrograms for two different subtypes of alarm calls, categorized into distinct clusters by HDBSCAN: clusters cl. 0 and cl. 10.

More detailed statistical analyses of the clusters and whether they are best explained by call type, identity or characteristics of the caller, or other variables are beyond the scope of this work but provide an interesting opportunity for studying the encoding of information in vocal signals.

#### Validation of the latent space representations

To compare the latent space representations of spectrograms with those generated from the extraction of acoustic features, we extracted 99% energy duration, cepstral peak prominence, centroid frequency, peak frequency, root mean square (RMS) bandwidth, fundamental frequency (F0) mean, F0 start, F0 mid, and F0 end from the meerkat calls. An additional variable ΔF0 was generated to capture the change in fundamental frequency over time by subtracting F0 start from F0 end. Signal-to-noise ratio (SNR) was not included in the analysis but was computed as a filtering step to discard weak calls (SNR<10 dB, N=933) for which features could not be reliably extracted. Noise level for SNR was calculated using the minimum of the lowest RMS noise level in a 100 ms window either preceding or following the signal. The remaining dataset of N=5,495 calls represented by the eight acoustic features was *z*-score normalized across features and projected into 3D space using *umap-learn* [7] with default hyperparameters except *min_dist=0*. For comparison, we also used the SNR-filtered dataset (N=5,495) to generate the spectrogram-based embedding. Both embeddings were then evaluated based on k=5 nearest neighbors and silhouette scores of manual call type labels.

Overall, the evaluation matrices of the spectrogram-based and acoustic feature-based embeddings were very similar. However, the embedding quality was higher for the spectrogram (*S*=63.38, *S*_*norm*_=2.37) than for the acoustic feature approach (*S*=52.20, *S*_*norm*_=2.06) (Figure 12).

**Figure 12:**
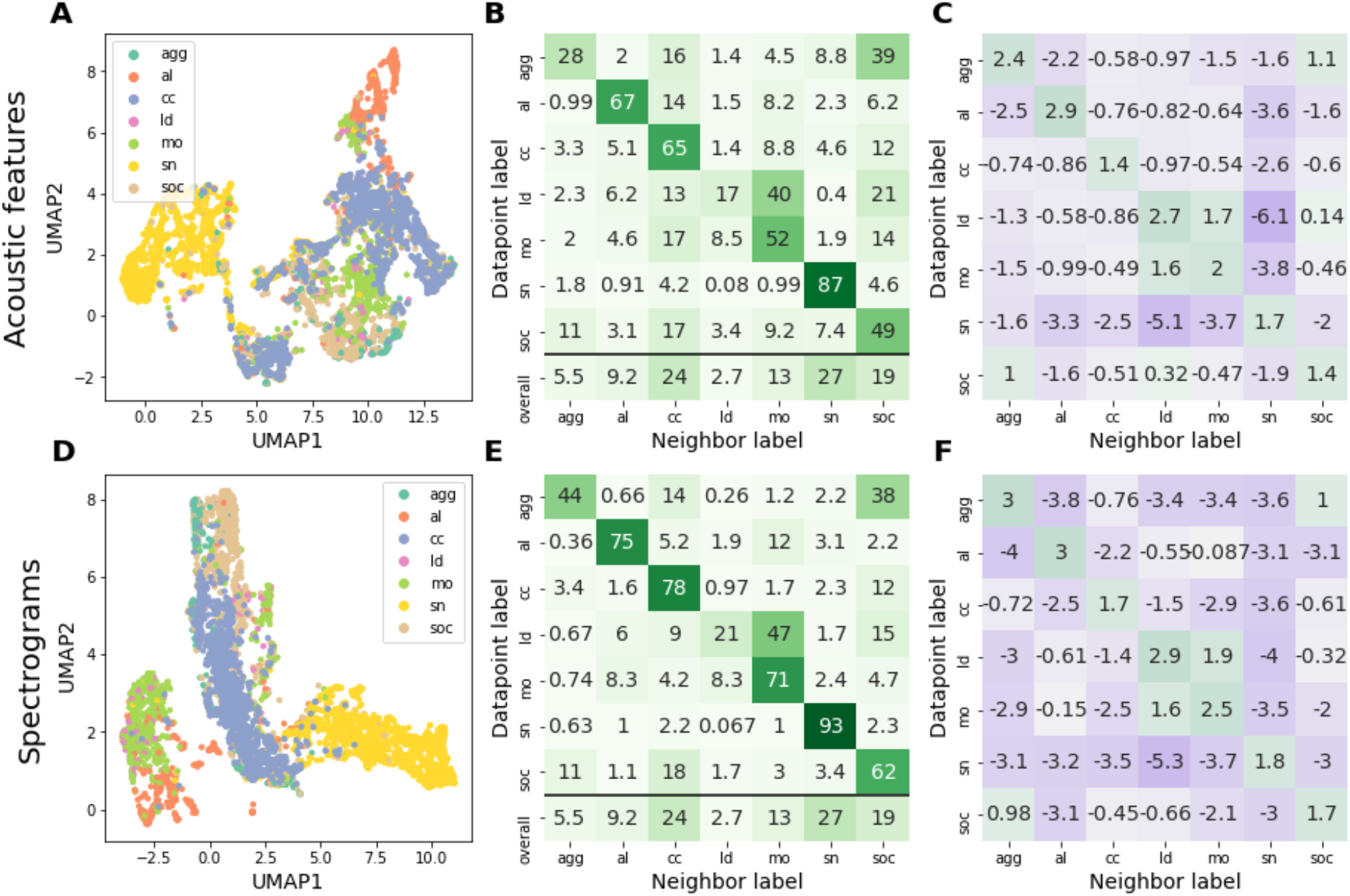
Comparison of UMAP embeddings generated with eight acoustic features (A-C) and with spectrograms as input (D-F). A) and D) are visualizations in 2D UMAP space. Evaluation matrices are based on k=5 nearest neighbors in 3D UMAP space. B) and E) show the absolute probability (in percentage) of encountering a neighbor with label y within the nearest neighbors of a datapoint with label x. C) and F) display log2-transformed ratio of that probability and the probability of encountering the neighbor label by chance. Analyses were performed with the reduced dataset, filtered for calls with SNR>10 dB (N=5,495). Colors are mapped on a violet-green scale from minimum to maximum value.

When comparing the silhouette values of the manual label classes, the SIL (average over all datapoints) was higher in the spectrogram-based vs. the acoustic feature-based embeddings (SIL=0.23 vs. SIL=0.13). The differences in silhouette values were most apparent for *mo* and *cc* calls, which formed better clusters in the spectrogram UMAP space than in the acoustic feature UMAP space (Figure 13).

**Figure 13:**
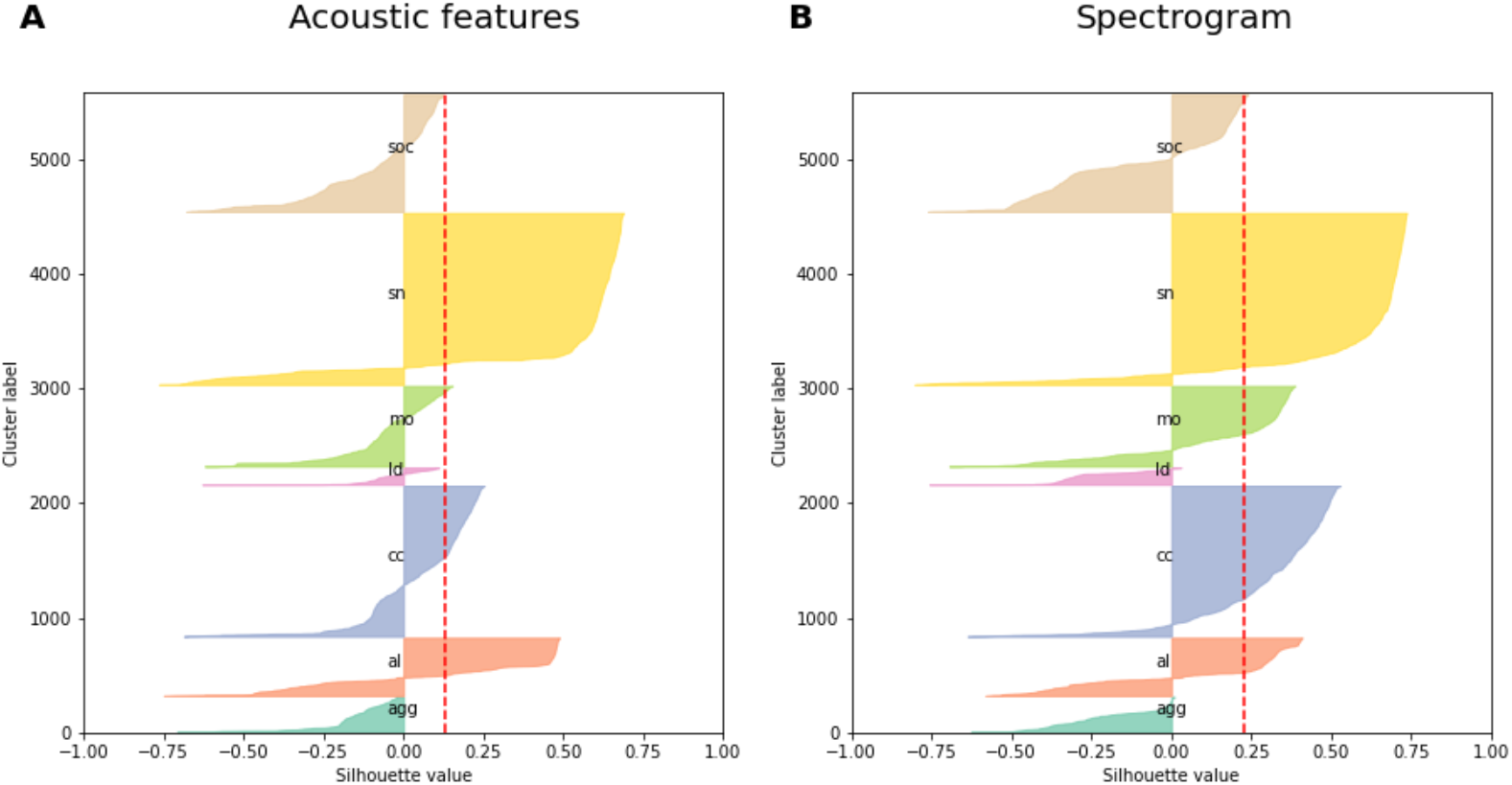
Silhouette plots for manual label classes in UMAP space generated with extracted acoustic features (A) and spectrograms (B). Silhouette values of all datapoints are represented as horizontal bars, grouped by call type label and sorted in descending order of silhouette values within each call type. Red dotted line shows average of all datapoints (SIL).

In summary, manually selecting and extracting specific acoustic features based on expert knowledge did not lead to a better local or global clustering of call types in the feature space than the simpler and less labor-intensive approach of using the spectrograms as feature vectors.

### TOOLS

The analyses presented here require a running installation of Python 3.8., Jupyter notebook and various core packages (Table 1):

**Table 1:** Overview of the core packages needed for the analysis.

| Category | Name | Version | Purpose |
| --- | --- | --- | --- |
| Basic computational pipeline | pandas | 1.2.4 | Data handling |
|  | numpy | 1.20.1 | Data handling, scientific computing |
|  | pysoundfile | 0.10.3 | Reading audio data |
|  | librosa | 0.8.0 | Generating spectrograms |
|  | umap-learn | 0.5.1 | UMAP |
|  | HDBSCAN | 0.8.27 | Clustering |
| Latent space evaluation<br>and adaptations of the<br>basic pipeline | scikit-learn | 0.24.1 | Nearest neighbor search, silhouette score |
|  | pygraphviz | 1.3 | Neighborhood graph |
|  | networkx | 2.5 | Neighborhood graph |
|  | numba | 0.53.1 | Custom distance function implementation |
| Visualization | matplotlib | 3.3.4 | Plotting |
|  | seaborn | 0.11.1 | Plotting |
|  | plotly | 4.14.3 | Interactive visualization |
|  | jupyter-notebook | 6.1.4 | Interactive visualization |
|  | ipython | 7.22.0 | Interactive visualization |
|  | ipywidgets | 7.5.1 | Interactive visualization |
|  | widgetsnbextension | 3.5.1 | Interactive visualization |
|  | voila | 0.10.2 | Interactive visualization |

For a full list of packages and dependencies, see conda environment file in supplementary materials.

### TRY-IT-YOURSELF

Our example dataset of 6,428 meerkat calls (.wav files), together with a tutorial-like set of jupyter notebook files to generate UMAP representations, the interactive visualization, HDBSCAN clustering and all presented evaluations from any set of input sound files are provided in a public github repository at https://github.com/marathomas/tutorial_repo and at https://doi.org/10.5281/zenodo.5767841.

## OTHER POSSIBILITIES AND DEVELOPMENTS

Even though the exemplary use case for unsupervised dimensionality reduction is clustering for the sake of detecting structure in an unlabelled dataset, we found that the approach is also useful for several applications in labelled or partially labelled datasets.

### Call type neighborhood graph

To visualize the degree to which different call types are acoustically similar to one another, we constructed a neighborhood graph where nodes represent call types and edges represent the probability of finding the connected call types among the k=5 nearest neighbors in latent space (Figure 16). In more detail, we transformed the evaluation matrix of the normalized k=5 nearest neighbor probabilities (*P*_*norm*_) of the 3D UMAP embedding into a symmetric distance matrix, replaced each field with the average of itself and its diagonal counterpart (*M*[*i,j*] = mean(*M*[*i,j*], *M*[*j,i*]) and then *M*[*j,i*]= *M*[*i,j*]), multiplied the matrix by -1 and set the diagonal to zero. We then generated an approximation of a graph where edge length represents the distance values from the matrix using *network* [21] and *pygraphviz* [22] (Figure 16). For the meerkat dataset, the resulting neighborhood relations were in line with the perceived similarity of call types by human listeners and, to a certain extent, with the function of calls (*mo* and *ld* calls are both associated with group movement, *agg* and *soc* calls are given primarily in social interactions).

**Figure 16:**
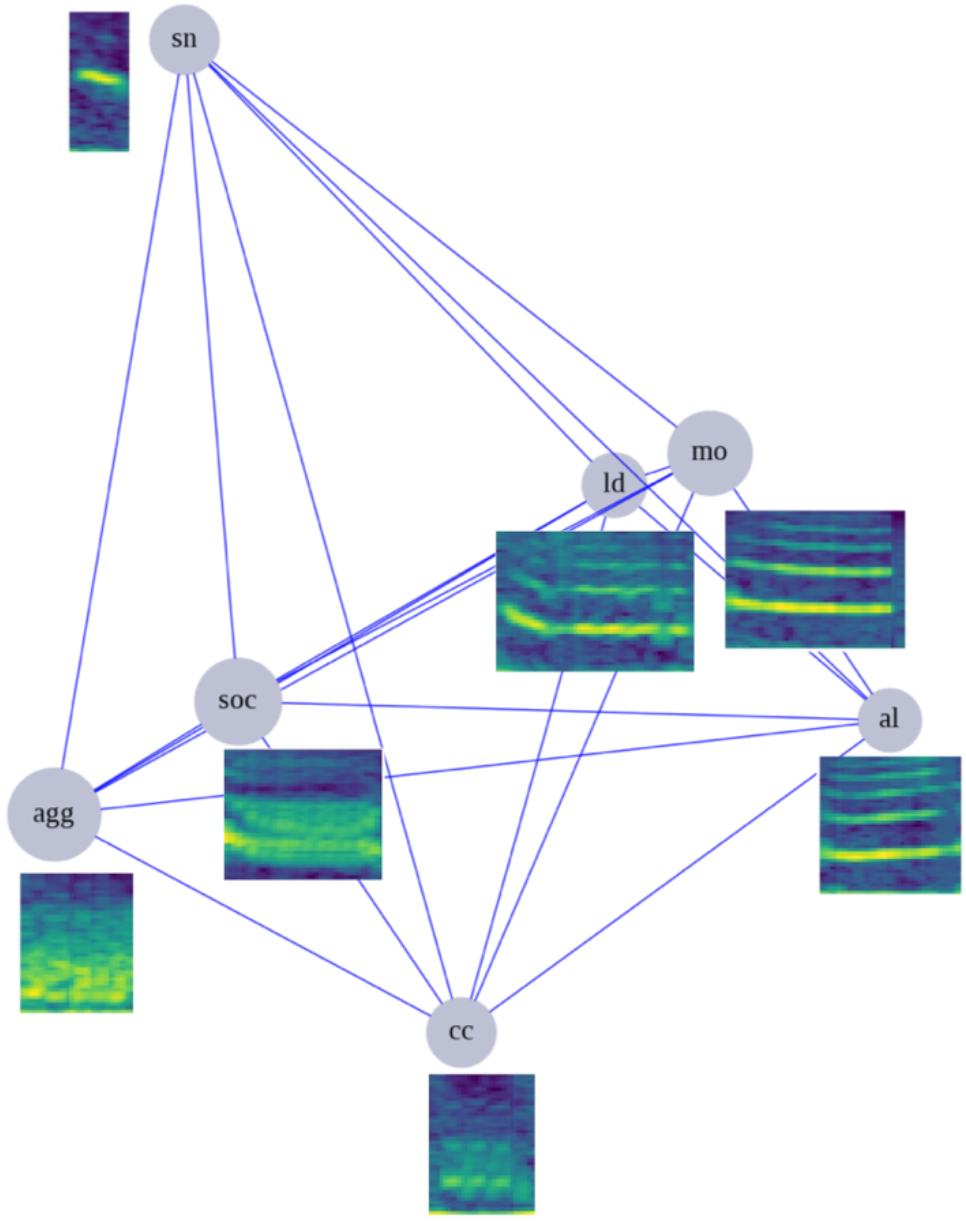
Call neighborhood graph based on k=5 nearest neighbors embedding evaluation. Call types connected by a shorter edge are more likely to be found in close vicinity to each other in the embedding. Example spectrograms are shown for each call type.

### Misclassification spotter

To test whether the local neighborhood in latent space can be used to identify mislabeled calls, we identified calls whose k=5 nearest neighbors all had a different label, randomly selected N=100 of these and asked two independent human experts to re-label the calls without providing any information on their previous assignment. 80 of the 100 calls were indeed re-labelled differently than their previous assignment by at least one of the labellers and thus seem to be truly mislabelled or ambiguous. Only for a small fraction of these (N=21), both labellers agreed on the new assignment (“clear cases”) and 19 of these (90.48%) would also have been correctly re-assigned based on majority vote among their k=5 nearest neighbors in UMAP space. For the remaining 59 calls, labellers either both agreed that these did not belong to any of the main call types (N=9 calls labelled as hybrids, noise or unknown) or disagreed on their assignment (N=50 calls), indicating that these calls are atypical and difficult to classify. For 12 of the 20 remaining “false alarms” (neighborhood in UMAP space indicated mislabeling, but both labellers agreed that their previous assignment had actually been correct), the labelers’ comments indicated some degree of uncertainty about their assignment. In conclusion, while simply re-assigning call type labels based on nearest neighbor classification will likely introduce errors in the dataset, the local neighborhood of calls can be used to identify groups of calls with a high probability of misclassification error and thus speed up the process of error detection and elimination in large datasets.

### Classification of ambiguous calls

To test the usefulness of the latent space representations for the classification of ambiguous calls, we used N=737 vocal elements that had previously been excluded from the analysis because they could not be clearly assigned to a call type and had been labelled as hybrids between two types. We projected these ambiguous calls into the existing UMAP space using the *transform* function of *umap-learn* [7] and assessed whether their k=5 nearest neighbors matched both or any of the call types from which they were presumably composed.

The projected hybrid calls were, as expected, distributed across the entire latent space, as opposed to forming their own cluster (Figure 17A). In most cases, the average percentages of call types in the neighborhood of hybrid calls were higher for those call types from which the hybrid call was composed (Figure 17B). Neighbors were also more likely to be acoustically similar to one of the hybrid labels. For example, *cc* calls were present among the nearest neighbors of *hyb:soc_agg* hybrid calls, and are acoustically similar to both types. When assigning a label to each hybrid call based on the majority call type among its nearest neighbors, 72.0% of calls were assigned to one of the designated hybrid labels, 25.8% to a completely different call type and 1.6% did not have a majority fraction (tie). When visually inspecting the 26.3% presumably mispositioned hybrid calls, the similarity between them and their nearest neighbors was evident (see supplementary information P4). Thus, a majority vote against any of the hybrid labels does not necessarily mean the method has failed to position this call but can also mean that mislabeled calls in the dataset confound results. Since the quality of nearest neighbor-based classification of novel calls depends on the quality of the labelled dataset, it is advisable to use a subset of the data that contains only typical representatives of the classes, and/or set a threshold that only allows high-confidence assignments (e.g. 100% same-class neighbors). It is also important to keep in mind that calls that are generally overrepresented in the dataset are more likely to be among the k nearest neighbors of a hybrid call by chance alone and may thus bias the classification. However, since the local neighborhood of data points is non-random, a normalization to the random-chance expectation would likely introduce more bias instead of reducing it.

**Figure 17:**
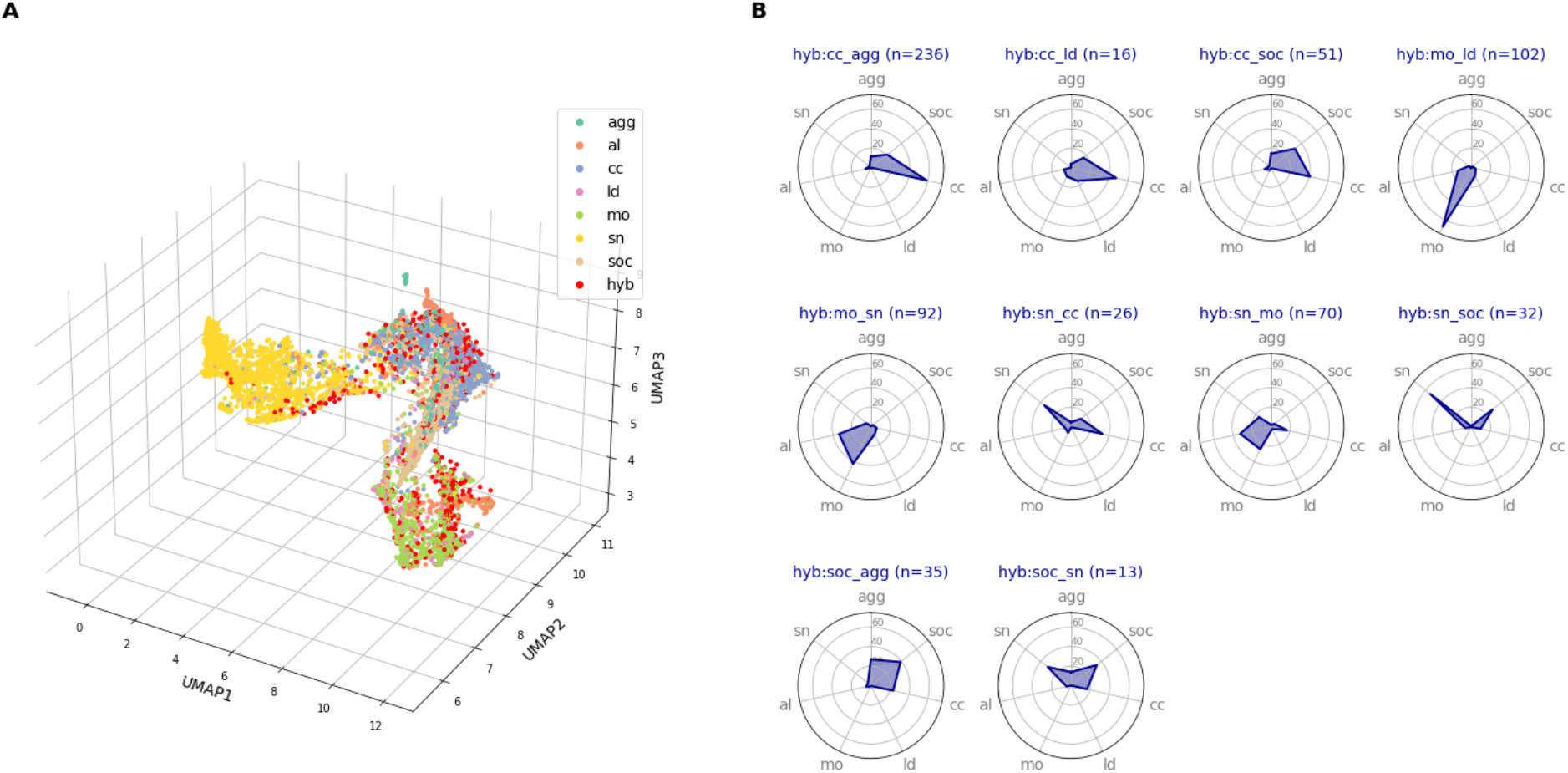
Classification of hybrid calls based on k=5 nearest neighbors. A) shows 3D UMAP plus hybrid calls in red (N=993). B) shows average percentage of call types among k=5 nearest neighbors of hybrid calls. The edges of the blue shapes represent the average percentages of different call types present among the k=5 nearest neighbors of hybrid calls. The order of call types on these charts was selected such that similar call types are next to each other. Chart titles indicate the two call types that the hybrid call is presumably composed of. Data is only shown for hybrid labels with N>10.

It is important to note that all of the presented applications could also be performed on the high-dimensional dataset. In fact, this will provide better results for small k, as the nearest neighbors in original space have slightly higher same-class neighbor frequencies for the local neighborhood (k<25, Figure 6). Strictly speaking, these applications thus do not demonstrate the usefulness of UMAP, but of the idea of assessing vocalization similarity by computing the distance between spectrogram-vectors. However, the differences between original and UMAP space are very minor and the use of UMAP space, while being less accurate, has the advantage that all steps can be visualized. In line with this, others have chosen to use UMAP for visualization, but performed computations in high-dimensional space [23].

## CAVEATS AND PITFALLS

### Meaningfulness of the distance metric

Altogether, the limitations of the latent space representations in visualizing acoustic similarity across a vocal repertoire are comprehensible considering how these representations were constructed, i.e. that the approach is based on element-wise comparison of spectrograms, similar to spectrogram cross-correlation [5]. This explains why certain types of signals appear similar in UMAP despite being perceived as acoustically distinct by animals and vice versa. For example, calls that are similar in shape, duration and tonality, but differ in continuity of the signal (e.g. pulsed vs. continuous signals in lead vs. move calls), will likely not be differentiated well, because they differ in only a few positions or elements of the spectrogram. In contrast, two signals of the exact same shape, but different intensity/loudness, would appear distant if the spectrograms were not normalized, which is why Decibel transformation and/or *z*-transformation are crucial preprocessing steps. Calls that have very similar shape but differ in duration or onset (especially calls with frequency modulation), calls with the same shape that are slightly different in frequency, calls with differences in the level and type of background noise or with different signal intensity can also appear very distant, even though they may in fact be acoustically similar. However, these issues are easy to anticipate and can be mitigated to some extent, e.g. by using dynamic time warping distance as distance metric or stretching all calls to the same length to account for differences in duration, by sliding spectrograms over one another in order to find the overlap position with minimum error (timeshift) and by using fewer Mel or frequency bins to alleviate the separation of calls with small shifts in frequency. Background and impulse noise can be attenuated, and several transformations can account for varying signal intensity between recordings. While some of these adaptations require more effort (e.g. custom distance metrics for *umap* may need to be implemented), others can easily be added to the pipeline (e.g. denoising or different types of normalizations) and our provided code contains implementations of many of these adaptations. Notably, all measures need to be carefully considered with regard to the aims of the analysis and the peculiarities of the dataset. For example, stretching or warping of calls should not be performed or be strictly constrained if duration is a biologically relevant acoustic feature. Similarly, tolerance limits for shifts in frequency depend on the vocal repertoire and the hearing abilities of the species of interest.

### Local and global structure preservation

Since UMAP favors preservation of local over global structure and is not designed to preserve all pairwise distances, distance in latent space cannot be interpreted as a proxy for distance in original space, especially for moderate to large distances. Further, even the local neighborhood in high dimensional space (i.e. the nearest neighbors) is not preserved exactly by UMAP (see supplementary information P5) [7]. Hence, while the projection to low dimensional space aids the visual and computational detection of clusters, many other downstream analyses (e.g. KNN classification) are more suitable to be performed in original, high dimensional space (as done in [23]). However, preservation of global structure can be increased if desired, either by increasing the number of neighbors for UMAP’s graph construction (*n_neighbors*), or by using Parametric UMAP [24], a recent addition to UMAP that enables the variation of global and local structure preservation through a parametric balancing between UMAP loss and an additional global structure preservation loss function.

### Bias imposition through parameter tuning

The freedom in choosing hyperparameters for generating and preprocessing spectrograms, as well as for running UMAP provides both the opportunity and the risk of tuning the analysis to the researcher’s needs. Assumptions about the relevance of specific properties of the signal (e.g. loudness or duration) will (and should) determine how spectrograms are generated, preprocessed and compared and thus define how acoustic similarity is assessed. It is important to be wary of this fact and not make the mistake of thinking that this (or any other) computational method can reveal “truth” in the data without a user having defined what aspects constitute this “truth”. In general, we recommend selecting all parameters prior to analysis and analyzing the effect of specific parameters on the outcome. However, we found that our analysis was very robust to changes in preprocessing and run hyperparameters overall (see supplementary information P7).

## ADDITIONAL PRACTICAL RESOURCES

In addition to our provided scripts, we recommend the original publication of the method [3] and the respective github repository: https://github.com/timsainb/avgn_paper. For information on UMAP beyond the original publication [7], we recommend the official documentation (https://umap-learn.readthedocs.io/en/latest/index.html), as well as this tutorial by Andy Coenen and Adam Pearce (Google People + AI Research): https://pair-code.github.io/understanding-umap/.

## CONCLUSION

Altogether, UMAP of spectrograms can produce meaningful representations of vocalizations, which are not inferior to those generated from commonly used acoustic features and are useful for a range of downstream applications beyond visualization and clustering of call types. Due to the speed and simplicity of the approach, it can also be useful for quality control in large datasets, automated classification, as well as for answering biological questions. We recommend fine-tuning the general framework of this computational approach to the needs of the specific analysis, while being wary of imposing bias to the analysis.

## Supporting information

Visualization tool demo video

## AUTHOR CONTRIBUTIONS

ASP, BA, and VD collected the data with support from MBM. BA, VD, and ASP manually annotated the acoustic data with input from MBM and with help from research assistants. MT carried out all analyses presented here with the exception of acoustic feature extraction, which was performed by FHJ. ASP, MAR, FHJ and TS provided scientific input and guidance on the analyses. MT wrote the tutorial code and the original draft of the manuscript. ASP tested the tutorial code. ASP and BA performed the manual re-evaluation of potentially mislabeled calls. All authors contributed to the manuscript and gave final approval for publication.

## ACKNOWLEDGEMENTS

We are grateful to the Kalahari Research Trust and Tim Clutton-Brock for the permission to work at the Kuruman River Reserve research site. We thank the Northern Cape Conservation Authority for research permission (FAUNA 1020/2016) and the managers and volunteers of the Kalahari Meerkat Project (KMP). This work was supported by HFSP Research Grant RGP0051/2019 to ASP, MBM, and MAR, and funded by the Deutsche Forschungsgemeinschaft (DFG) under Germany’s Excellence Strategy (EXC-2117-422037984). ASP received additional funding from the Gips-Schüle Stiftung, the Zukunftskolleg at the University of Konstanz, and the Max-Planck-Institute of Animal Behavior. VD was funded by the Minerva Stiftung and Alexander von Humboldt Foundation. We thank Gabriella Gall and Rebecca Schaefer for assistance with field work, as well as Richard Young, Silvan Spiess and Leonardos Leonardos for assistance with call labeling.

