## Supplementary information for "A practical guide for generating unsupervised, spectrogram-based latent space representations of animal vocalizations"

### P1: Data collection and preprocessing

The original data consists of 108 hours of audio recordings of wild meerkats. These data were collected during two field seasons (Aug-Sep 2017 and Jul-Aug 2019) at the Kalahari Research Centre in South Africa. Audio was recorded with acoustic collars (TS-market Edic Mini Tiny+ A77, samplerate 8,000 Hz) that were attached to the animals, as well as with portable digital recorders (Marantz PMD661) and directional microphones (Sennheiser ME66) (both sample rate 48,000 Hz), that field researchers held close to the animals (within 1 m). 29,569 meerkat calls were identified in these recordings by human listeners and their start and stop positions marked using Adobe Audition. This number excludes calls that were labelled as noisy, that could not clearly be identified, that could not be categorized into any of the seven major call types or that were non-focal calls, i.e. produced by animals in the background and not by the animal wearing the acoustic collar. The identified calls had been labeled as either aggression (*agg*), alarm (*al*), close (*cc*), lead (*ld*), move (*mo*), short note (*sn*) or social call (*soc*). All calls are continuous vocalizations that are preceded and followed by silence, yet no strict threshold for the minimum duration of the silent period in between calls was applied. Consecutive vocalizations were categorized as separate calls if the labeler perceived them as distinct. The duration of calls was approximately normally distributed around a mean of 135 ms. To obtain a clean and more homogeneous sample, calls longer than 500 ms (282 calls, 1.3 % of the dataset) and shorter than 50 ms (2,137 calls, 9.8 % of the dataset) were removed from the dataset (N=19,722 remained). To attenuate the overrepresentation of close calls in the dataset without losing any other calls in the dataset, we reduced the number of close calls in the dataset to a random 10 % subsample (from N=14,769 to N=1,477). Six duplicate calls were removed from the dataset, leaving N=19,716 calls in the full and N=6,428 in the reduced dataset.

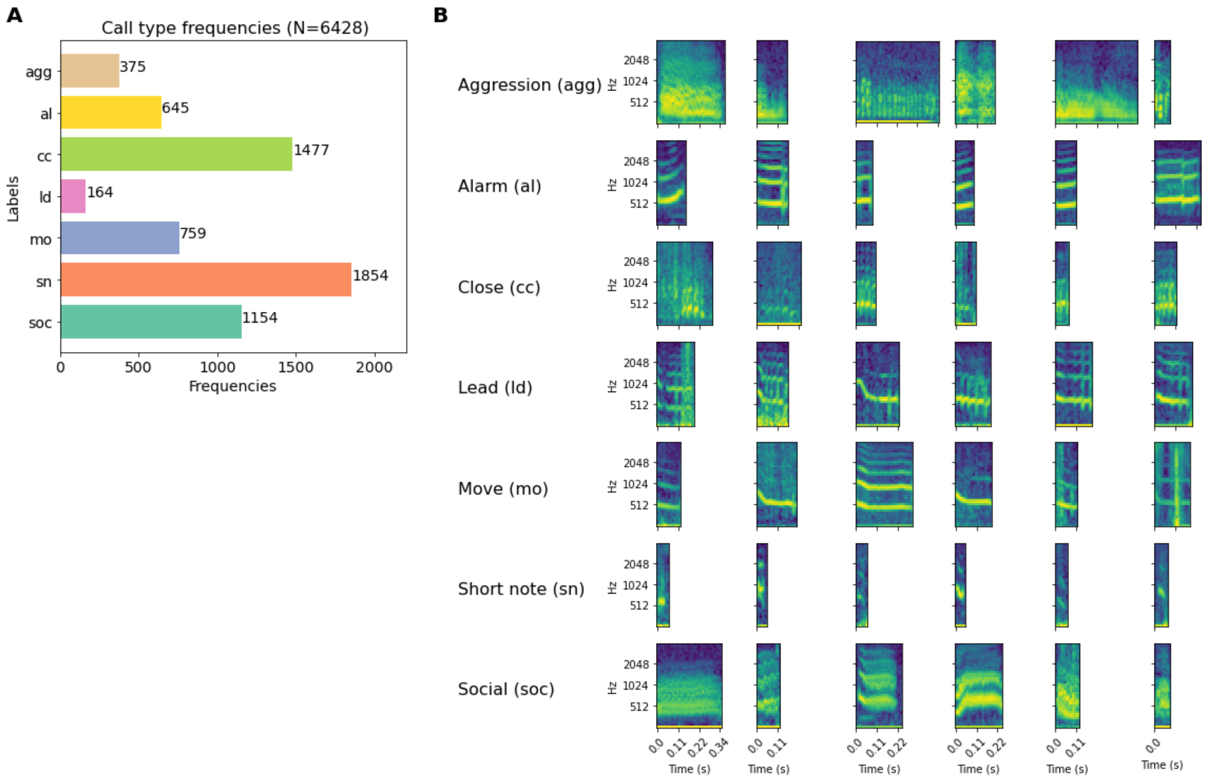


Figure S1: A) Absolute frequencies and B) example spectrograms of the different call types in the dataset

### P2: Effect of class imbalance

To test, whether the reduction of the overrepresented *cc* fraction or class imbalances in general had a substantial effect on our results, we generated UMAP embeddings with the full dataset, as well as a completely balanced one, where we downsampled each call type to the rarest class (lead calls N=164, total N=1,148). All embeddings were evaluated using the nearest neighbor-based evaluation scores $S$ and $S_{norm}$.

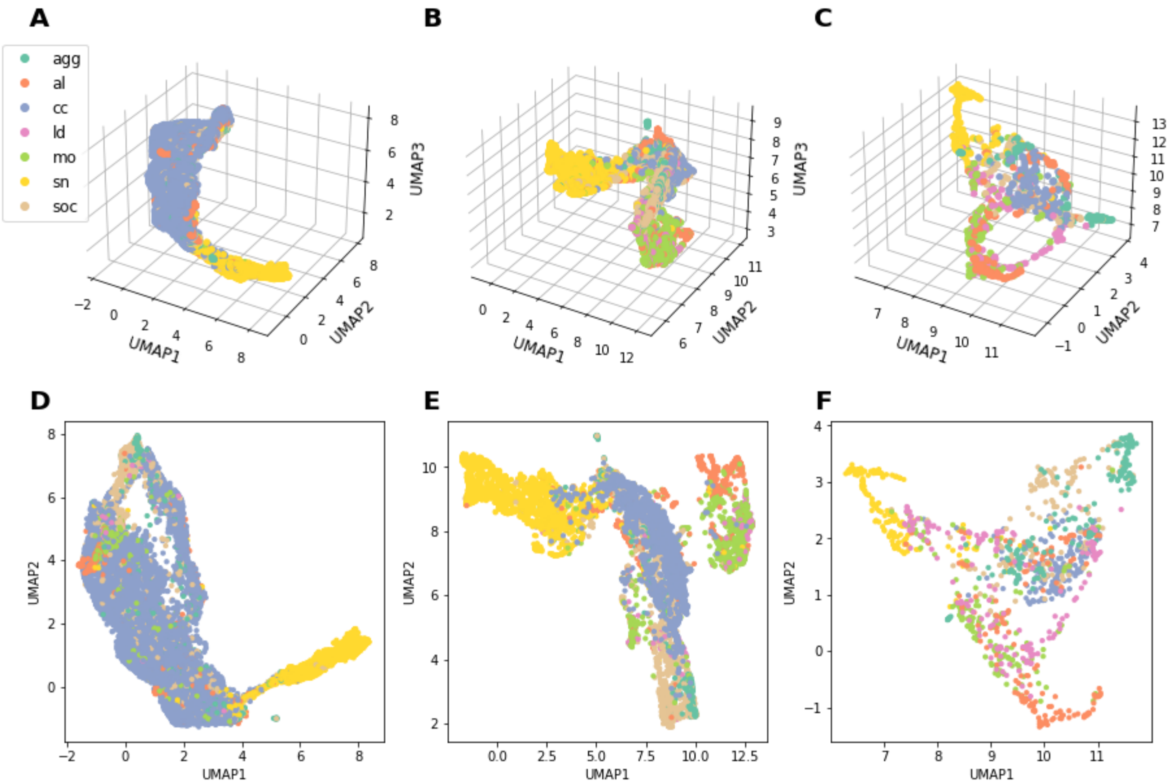


*Figure S2: Meerkat call embeddings in 2D (A-C) and 3D (D-F) UMAP space, created using row-wise concatenated, zero-padded spectrograms as input. Shown for the full, reduced and balanced dataset. Datapoints are color-coded by manual labels.*

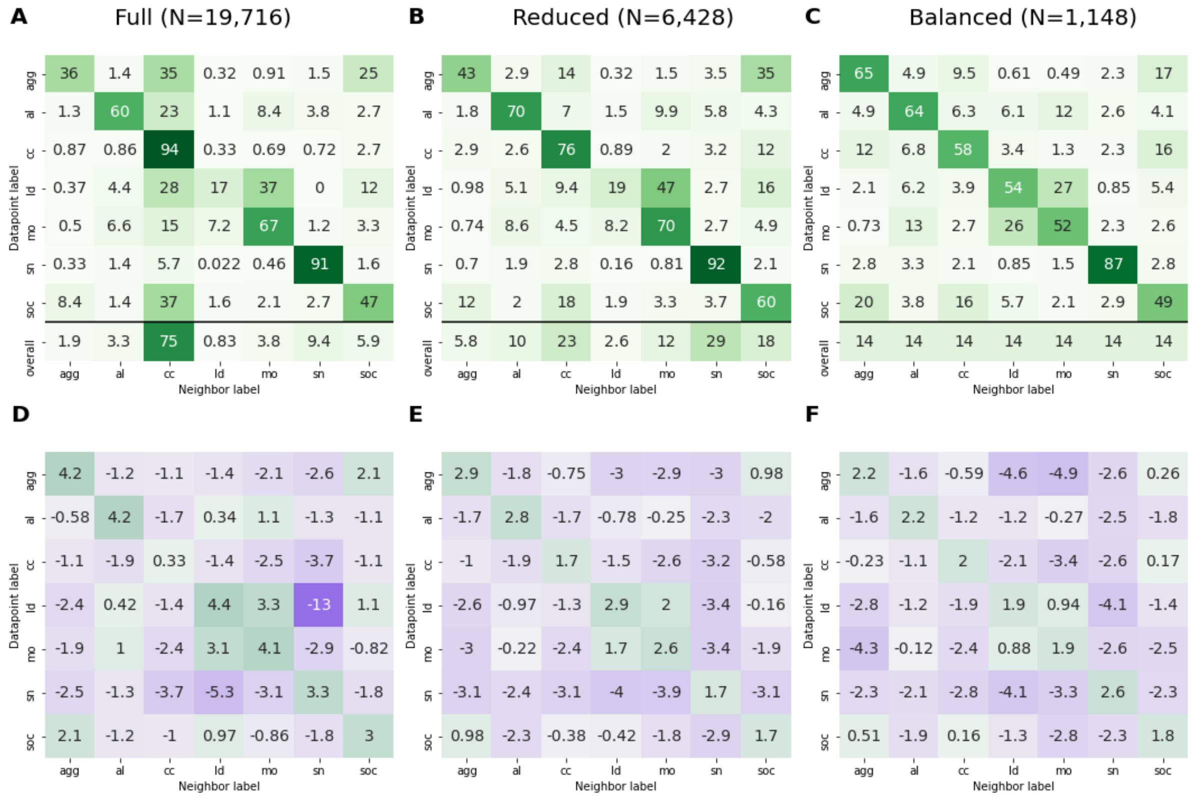


*Figure S3: Evaluation metrics for 3D UMAP embedding of full, reduced and balanced datasets. Calculations are based on evaluation of k=5 nearest neighbor in 3D space. Confusion matrices A)-C) show the absolute probability (in percentage) of encountering a neighbor with label y within the k=5 nearest neighbors of a datapoint with label x. D)-F) display log2-transformed ratio of that probability and the probability of encountering the neighbor label by chance (i.e. a normalization to label frequencies in the dataset). Colors are mapped on a yellow-red scale ranging from minimum to maximum value.*

Overall, the quality score $S$ is highest for the balanced (S=61.36), followed by the reduced (S=61.35) and full dataset (S=58.74). The quality score $S_{norm}$ is highest for the full dataset ($S_{norm}$=3.36), followed by the reduced ($S_{norm}$=2.33) and balanced ($S_{norm}$=2.08) dataset. The incongruent rank order for $S$ and $S_{norm}$ is caused by the unequal class frequencies in the three datasets (see lowest row of evaluation matrices in Figure S3A-C). Since $S_{norm}$ is normalized for these class frequencies, it is better suited to compare datasets of varying composition. However, $S_{norm}$ (the unweighted mean of $P_{norm}$ over all call types) needs to be interpreted with caution. For example, the strong overrepresentation of a specific call type causes low same-class $P_{norm}$ for this type, and high same-class $P_{norm}$ for all other call types, even if the absolute neighbor probabilities $P$ may indicate otherwise. This is apparent in the full dataset, where close calls represent 75 % of the dataset. The normalized same-class score is very low for close calls ($P_{norm}$=0.33), even though the absolute neighbor probability is very high at 94 % (Figure S3 A, D). Hence, we recommend assessing both normalized and unnormalized evaluation metrics for evaluating the quality of an embedding.

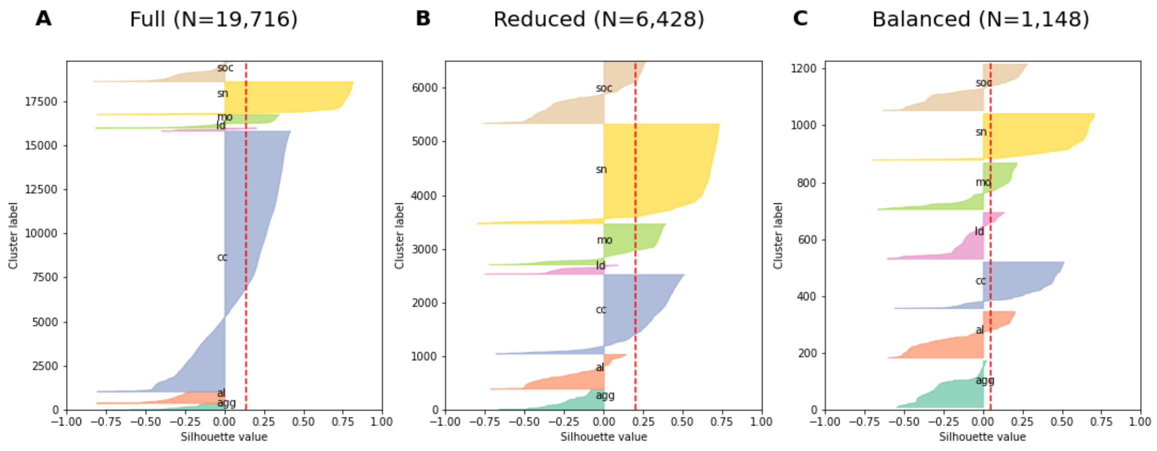


*Figure S4: Silhouette scores for manual label clusters in 3D UMAP embeddings of A) full, B) reduced and C) balanced datasets. Datapoints are sorted by label and each silhouette value for each datapoint is displayed. Dotted red line indicates the average silhouette score for all datapoints.*

When looking at average silhouette values for the manual call type groups, these were highest for the reduced (SIL=0.2), followed by the full (SIL=0.15) and balanced (SIL=0.05) dataset. This is likely caused by the high overrepresentation of well-clustered *cc* calls in the full dataset. When weighing each call type equally in the average silhouette value (as is done for $S$ and $S_{norm}$), the reduced and balanced dataset have similar scores (SIL=0.05 for both), whereas the full dataset has SIL=0.01. Thus, heavy overrepresentation of particular call types may negatively affect clusterability, however class imbalances do not alter the overall patterns of local neighborhood in latent space.

### P3: Effect of dataset size

To assess the effect of dataset size, we computed UMAP embeddings and assessed the k=5 nearest neighbor probabilities for datasets with varying numbers of samples per class, while keeping the class frequencies constant. To avoid effects of class imbalances, we used the completely balanced dataset for this analysis.

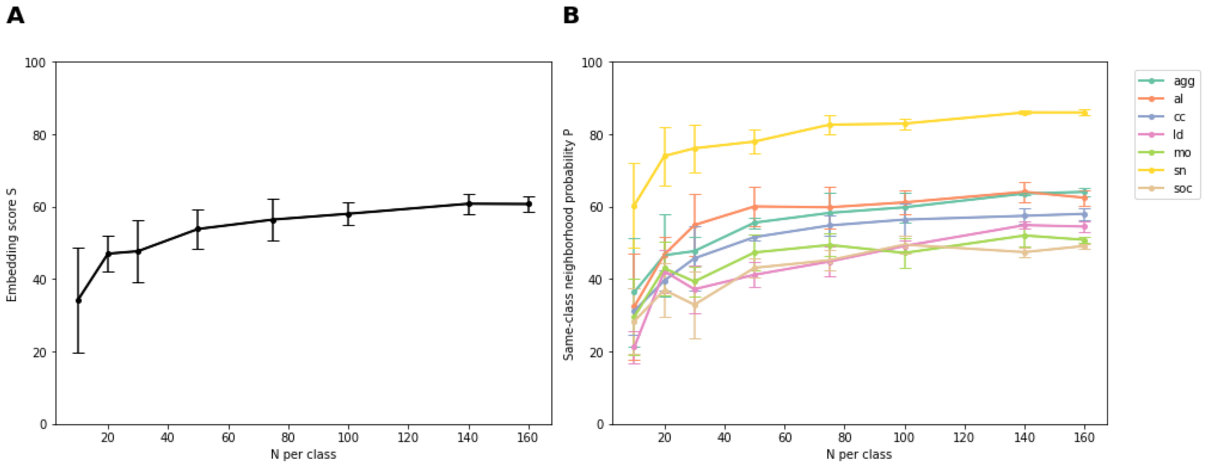


*Figure S5: Embedding scores S (A) and same-class neighborhood probability P (B) for embeddings generated with differently sized subsets of the balanced dataset (number of samples per class ranged between N=10 and N=164). Datapoints are means plus standard deviations from n=5 random subset samplings.*

The quality scores $S$ and $S_{norm}$ increase with the number of calls per class in the dataset, but the curve flattens above approximately N=50 calls per class (Figure S5). Short note calls are the class with the highest same-class neighborhood probability for all dataset sizes and the rank order of all other calls also remains mostly stable from N=30 calls upwards. The variance in quality scores is high for datasets with N<30 samples per class and smaller for all datasets with N>=30. Altogether, these results indicate that N=50 calls per call type are sufficient to achieve similar quality scores of same-class nearest neighbors as in the complete dataset with N=191 calls per call type. While these values certainly depend on the dataset and choice of k nearest neighbors and may thus vary for other datasets, they still provide an orientation for a minimum number of vocalizations that are necessary explore a vocal repertoire based on the nearest neighbors in UMAP space.

### P4: Inspection of mispositioned hybrid calls

When projecting hybrid calls to latent space and assigning a class label based on majority vote among the k=5 nearest neighbors, 26.3 % of the calls were assigned to a call type that was not one of the designated hybrid labels. However, visual inspection of these presumably mispositioned hybrid calls showed the similarity between them and their nearest neighbors.

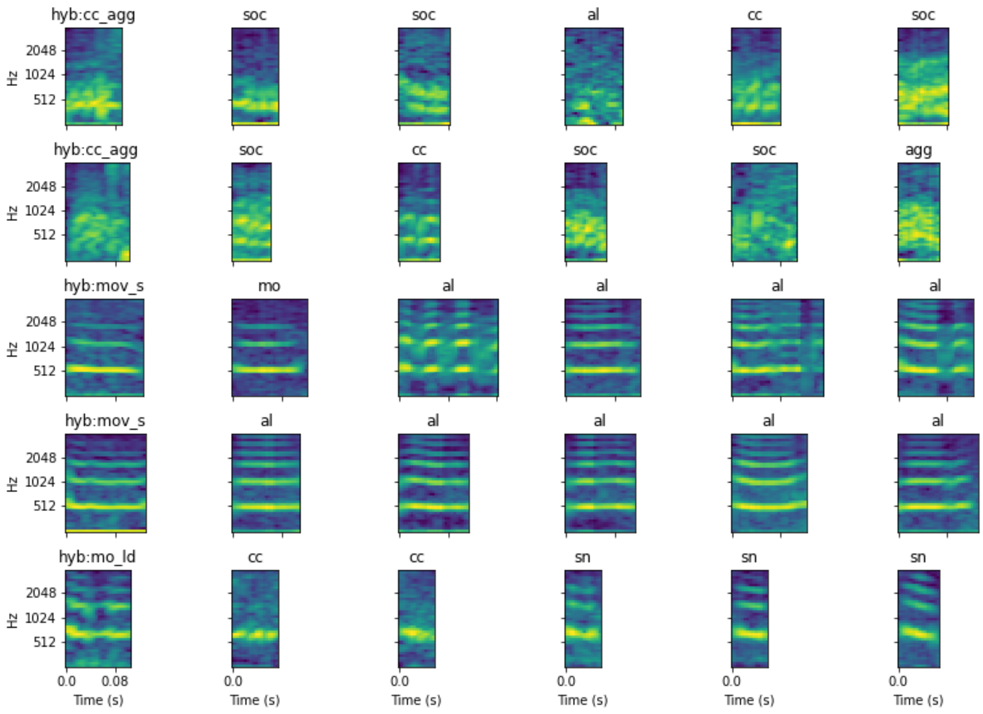


*Figure S6: N=5 randomly pulled example hybrid calls, where the majority vote among the k=5 nearest neighbors in latent space was none of the designated hybrid label call types. The hybrid call are shown in the left column and their k=5 nearest neighbors in columns 1-5. Titles indicate manually annotated call type label.*

### P5: Computation of nearest neighbor preservation

To assess how well local structure is retained in the low dimensional representation, we identified the k=5 nearest neighbors in low- and high-dimensional space (e.g. UMAP vs. original) and calculated the percentage of agreement between the two, e.g. the fraction of k=5 nearest neighbors in original space that were also among the k=5 nearest neighbors in UMAP space. The nearest neighbor preservation was 26.1 %, indicating a low exact preservation of neighbors. This is most likely due to the high dimensionality of the spectrogram-vectors (>5,000 dimensions), making it impossible to model these accurately in low dimensional space. In addition, it may be exacerbated by inaccuracies in the nearest neighbor approximation of UMAP and the fact that UMAP constructs its neighborhood graph based on neighborhood probabilities, which do not depend on distance alone, but also take local density into account.

### P6: Correlation of distances in low- and high-dimensional space

To assess whether distances in original space are similarly modeled in UMAP space, we calculated the Pearson correlation coefficient of the distance matrices in original and UMAP space using the Mantel test implemented in *scikit-bio* v0.5.0 [1] with default parameters (n=999 permutations). In addition, we grouped all pairs of points into 50 equal-width bins based on their distance in the original space. For each bin, we plotted the distance for the pairs of points in the embedded space, recreating a figure from Becht et al. [2] to visualize preservation of distances in low dimensional space.

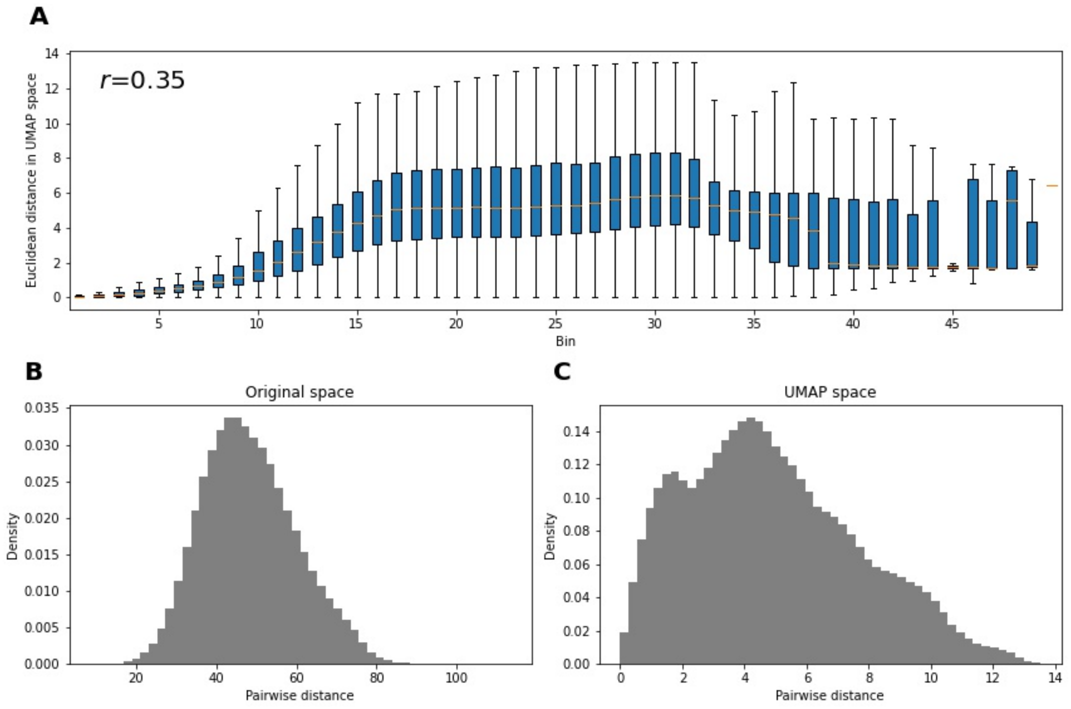


*Figure S7: Preservation of pairwise distances in embedding. A) An analogue of Figure 5 in Becht et al. [2]. Quoting the original caption: "Box plots represent distances across pairs of points in the embeddings, binned using 50 equal-width bins over the pairwise distances in the original space [...] The value of the Pearson correlation coefficient computed over the pairs of pairwise distances is reported. For the box plots, the central bar represents the median, and the top and bottom boundary of the boxes represent the 75th and 25th percentiles, respectively. The whiskers represent 1.5 times the interquartile range above (or, respectively, below) the top (or, respectively, bottom) box boundary, truncated to the data range if applicable." Datapoints outside of 1.5 times the interquartile range are not shown. B),C) show the histograms of pairwise distances in B) original and C) UMAP space.*

The Pearson correlation coefficient was low ($r$=0.35). Overall, distances in the embedded space increased along with the distances in the original space for small pairwise distances, remained largely constant for moderate distances and decreased with increasing distances in the original space. This behavior is expected, as the neighborhood probability in embedded space is modeled by a nearly constant, low value when the neighborhood probability between two points in original space is below a certain value (i.e. their distance is large). The visualization of the distance correlation emphasizes that moderate to large distances between datapoints in UMAP space should not be over-interpreted, as they may not actually reflect large distances in original space.

### P7: Adjustment of the UMAP processing pipeline

Numerous adjustments can be performed to tune the computational pipeline to meet the needs of specific analysis questions. To provide an overview of some possible adjustments, we evaluated the effect of different inputs, preprocessing steps and UMAP hyperparameters on the embedding score *S* of the latent space representations by performing an exhaustive grid search with a set of predefined hyperparameter values on our dataset. With regard to spectrogram generation and preprocessing, we tested different numbers of Mel coefficients (10, 20, 30, 50), as well as omitting the Mel-transformation, spectrograms with magnitude unit instead of Decibel, and spectrograms that had not been normalized by *z*-transformation. To deal with different types of noise, we tested the effect of applying a floor and ceiling value to the *z*-transformed spectrograms (floor: 0, ceiling: 3), bandpass filtering of frequencies <300 Hz and >3 kHz to remove background noise and median subtraction per spectrogram column (STFT time frame) to attenuate the effect of impulse noise specifically. With regard to UMAP hyperparameters, we tested different values for UMAP spread (0.5, 0.75, 1.0 and 1.5), UMAP n_neighbors (5, 10, 15, 30, 50, 100, 150, 200), UMAP min_dist (0.0, 0.001, 0.01, 0.1 and 1.0) and UMAP distance metric (Correlation, Cosine, Euclidean, Manhattan). Lastly, we tested different ways of dealing with the varying duration of vocalizations. In addition to zero-padding of spectrograms to a maximal length as described in [3], we also stretched all spectrograms to the maximal call duration in the dataset (500 ms) using a phase vocoder algorithm [15] (stretch), zero-padded the spectrograms up to the length of the longer spectrogram of each pair that is compared (pairwise-pad), calculated the distance only from overlapping sections of the spectrogram that are aligned at the start (overlap), shifted the spectrograms in time to find the overlap position with the lowest distance and then calculated the distance only from the overlapping section (tshift-overlap) or zero-padded the smaller spectrogram to match the duration of the longer one (tshift-pad).

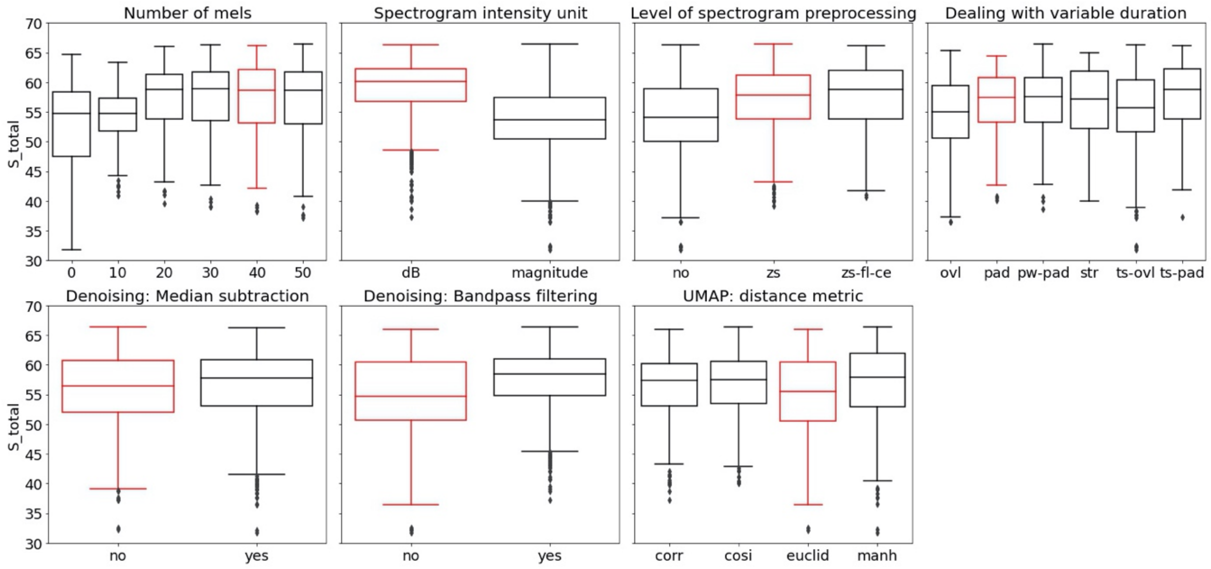


*Figure S8: Results of grid search for evaluating parameters that define how well distance in original space reflects call similarity. Boxplots are shown for distribution of embedding scores for all parameter variables and their levels. Parameters that are used in the basic pipeline are highlighted in red. Abbreviations: Ovl: overlap, pad: padded, pw-pad: pairwise-pad, str: stretched, ts-ovl: tshift-overlap, ts-pad: tshift-pad. Corr: correlation, cosi: cosine distance, euclid: Euclidean distance, manh: Manhattan distance.*

Overall, the 99% confidence intervals for the quality score *S* ranged between 40.4 – 65.3% for all embeddings. Thus, irrespective of preprocessing steps and hyperparameters, spectrogram-based UMAP grouped calls of the same type together to a degree far above random chance (random expectation: 14.7%). Important parameters that increased the quality of the embedding were the transformations of frequency to Mel (with no difference between 20, 30, 40 or 50 Mels) and of magnitude to Decibel (Figure 14). Further, normalization by *z*-transformation improved the embedding, whereas the addition of a floor and ceiling to the *z*-tranformed intensity values only had a minor positive effect. Interestingly, the more sophisticated ways of dealing with variable duration calls did not bring a major improvement to the overall embedding. This is likely due to the fact that calls of the same type were very similar in duration in our dataset and their start position was accurately labelled. Hence, the signals in the spectrograms were well aligned even without any shifting in time. However, these aspects will likely be different for other datasets. With regard to denoising, bandpass filtering improved the embedding and the application of median subtraction did so to a lesser extent. This may also be different for datasets that contain more or different types of noise. Lastly, the use of correlation, cosine distance and Manhattan distance (L1 norm) as distance metric produced slightly better embeddings than Euclidean distance.

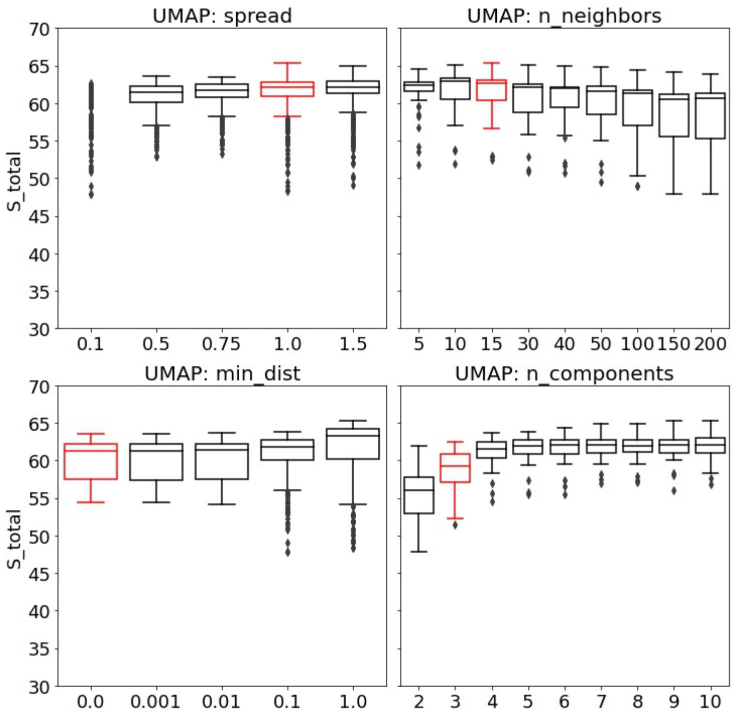


*Figure S9: Results of grid search for evaluating parameters that define how well distance in original space reflects call similarity. Boxplots are shown for distribution of embedding scores for all parameter variables and their levels. Parameters that are used in the basic pipeline are highlighted in red.*

Whereas most UMAP hyperparameters did not have a strong effect on the embedding score *S*, the number of components (i.e. dimensions of the resulting embedding) clearly improved the quality of the embedding up until four dimensions, after which it reached a plateau (Figure 15). Since all embeddings were evaluated with *S*, the unweighted average of the same-class *P* scores across all call types, it is possible that parameter combinations affected the clustering of the seven call types differently and this was not considered in the analysis. We also evaluated the average silhouette score for all embeddings and the direction of effects was identical except for *min_dist*, where *min_dist*=1 actually produced the lowest silhouette score. It is important to keep in mind that due to the large complexity, we did not consider interaction effects between different parameters here, even though these clearly exist (e.g. within UMAP hyperparameters *spread* and *min_dist* or levels of spectrogram preprocessing and denoising by median subtraction).
